# Dispersal increases the resilience of tropical savanna and forest distributions

**DOI:** 10.1101/476184

**Authors:** Nikunj Goel, Vishwesha Guttal, Simon A. Levin, Carla A. Staver

**Affiliations:** Department of Ecology and Evolutionary Biology, Yale University, New Haven, Connecticut, USA; Centre for Ecological Sciences, Indian Institute of Science, Bengaluru, Karnataka, India; Department of Ecology and Evolutionary Biology, Princeton University, Princeton, New Jersey, USA.

**Keywords:** dispersal, climate change, savanna-forest boundary, alternative stable states, resilience, source-sink dynamics.

## Abstract

Global change may induce changes in savanna and forest distributions, but the dynamics of these changes remain unclear. Classical biome theory suggests that climate is predictive of biome distributions, such that shifts will be continuous and reversible. This view, however, cannot explain a widely observed mismatch between climate and tree cover, which some argue results from fire-vegetation feedbacks maintaining savanna and forest as bistable states, such that, instead, shifts will be discontinuous and irreversible. This bistable model, however, cannot reproduce the spatial aggregation of biomes. Here, we suggest that both models are limited in that they ignore spatial processes, such as dispersal. We examine the contributions of dispersal to determining savanna and forest distributions using a reaction-diffusion model, comparing results qualitatively to empirical savanna and forest distributions in Africa. The diffusion model induces spatially aggregated distributions, separated by a stable savanna-forest boundary. The equilibrium position of that boundary depends not only on precipitation but also on the curvature of precipitation contours with some history dependence (although less than in the bistable model). This model predicts different dynamics in response to global change: the boundary continuously tracks climate, recovering following disturbances, unless remnant biome patches are too small.

## Introduction

Climate and land use change are expected to result in large-scale shifts in global vegetation patterns (Aleman et al. 2016; Loarie et al. 2009; Malcolm et al. 2002; Salazar et al. 2007), leading to loss of biodiversity and ecosystem services that are vital for human livelihoods (Daily 1997). However, biosphere responses to changing climate and land use are uncertain. This uncertainty stems from uncertainty in what determines global biome patterns; current biome distribution models are unable to explain even simple empirical features of today’s vegetation patterns. Predicting changes in biome distributions with respect to global change thus requires a better understanding of the drivers and the mechanisms by which these drivers shape global biome patterns.

Conceptually, the classical theory suggests that climate is the fundamental determinant of vegetation pattern and that there is a one-to-one match between climate and biome (Holdridge 1947; von Humboldt and Bonpland 1807; Whittaker 1970), such that biomes continuously track changes in climate through space. Thus, under the classical view, biome shifts are continuous and reversible, and as such, relatively predictable. An alternative viewpoint, supported by both field (Dantas et al. 2016) and remote-sensing approaches (Hirota et al. 2011; Staver et al. 2011a) suggests that a single climate can support multiple vegetation types, which are differentiated instead by other ecological processes including chronic fires (Bond et al. 2005; Staver et al. 2011a; Staver et al. 2011b). Savanna and forest may be a classic example; fire experiments have repeatedly shown that frequent fires can maintain savanna in regions where a closed canopy forest is climatically possible (Swaine et al. 1992; Trapnell 1959; Veenendaal et al. 2018). Simple theoretical models that incorporate both climate and fire suggest that savanna and forest may be bistable, with substantial hysteresis in biome patterns (Beckage et al. 2009; Staver et al. 2011b; Staver and Levin 2012). In this second scenario, unlike the classical theory, vegetation responses to changing climate and land use may be large and irreversible, and therefore difficult to foresee.

The bistable theory, however, has its limitations as well. Although the mean-field bistable models can mechanistically explain the overlap in the climatic ranges over which savanna and forest occur (Beckage et al. 2009; Staver and Levin 2012), they cannot be used to describe spatial patterning of biomes. Most notably, they miss obvious spatial features of savanna and forest distributions: savannas are found near other savannas and forests near other forests, with a distinct biogeographic boundary separating the two biomes (Aleman and Staver 2018). This spatial aggregation is not an obvious outcome of mean-field models, unless they invoke an additional assumption that the historical or paleo-distributions of savanna and forest are spatially structured by some extrinsic process (see Aleman and Staver 2018) (*e.g*., paleoclimate).

An alternative explanation could be that some spatial process at the savanna-forest ecotone may spatially aggregate savanna and forest. For instance, studies show that seed dispersal from forest patches can allow recovery of nearby derived savannas (Holl et al. 2000; Puyravaud et al. 1994) by clumping fragmented forest patches into bigger forest aggregates. Thus, dispersal could potentially explain the observed spatial aggregation of savanna and forest. However, only a handful of theoretical studies (Favier et al. 2004; van de Leemput et al. 2015; Wuyts et al. 2018) have explicitly considered the role of dispersal in determining biome patterns at relevant spatial scales.

The problem should be tractable, however, as dispersal is among the best-studied spatial ecological processes. Traditionally, dispersal in ecology is has been studied via reaction-diffusion equations. These equations offer a simple and analytically tractable way to incorporate dispersal in modeling dynamics of populations at large spatial scales (Levin 1992; Skellam 1951). Theoretical work on one-dimensional reaction-diffusion models shows that coupling diffusion with a bistable model (van de Leemput et al. 2015; Wuyts et al. 2018) can yield spatially aggregated biome distributions, separated by a stable savanna-forest boundary. Moreover, the one-dimensional reaction-diffusion model behaves, dynamically, like the classical biome theory: savanna and forest distributions continuously track changes in climate and recovers to equilibrium following perturbations. Unfortunately, this one-dimensional diffusion model (van de Leemput et al. 2015; Wuyts et al. 2018) also reverts to the main drawback of the classical biome theory: it, too, fails to reproduce the widely observed overlap in the climatic ranges over which savanna and forest biomes occur (Hirota t al. 2011; Staver et al. 2011a).

One obvious avenue for exploration is that these models that couple diffusion to models for savanna-forest dynamics treat the landscape as one-dimensional (1D) (van de Leemput et al. 2015; Wuyts et al. 2018), whereas, in reality, savanna and forest dynamics play out on two-dimensional (2D) landscapes. Going from 1D to 2D often gives rise to new dynamical features, such as motion by mean-curvature (Allen and Cahn 1979; Chen 1992; Evans et al. 1992; Gandhi et al. 1999; Keener 1986; Merriman et al. 1992; Tyson and Keener 1988), that could fundamentally change the dynamics of boundaries and thus their equilibrium distributions. Here, we address this directly by considering a model that couples the bistable mean-field vegetation structure with a diffusion process in two dimensions. In particular, we ask (*i*) whether seed dispersal, approximated as a two-dimensional diffusion process could contribute to spatial aggregation of savanna and forest biomes at continental scales; (*ii*) if yes, what then determines the equilibrium position of the savanna-forest boundary, and (*iii*) how this impacts the resilience of savanna and forest biomes to perturbations and global change. Finally, (*iv*) we empirically test some of the key analytical predictions of our 2D reaction-diffusion model using remotely sensed biome (Hansen et al. 2013) and climate patterns (Huffman and Bolvin 2013) in Sub-Saharan Africa.

### Model Description

Here, we present a reaction-diffusion model of savanna and forest biomes that consists of two parts: the reaction term that determines how fire interacts with vegetation and climate, and the diffusion term that represents seed dispersal. Here, we first describe the reaction term, and then, following, the diffusion term.

In savanna and forest ecosystems, fire exerts strong control over tree cover (Bond et al. 2003; Bond et al. 2005) via feedbacks with vegetation. In a low tree-cover landscape, fire spreads readily in the landscape (Archibald et al. 2009; Staver and Levin 2012) because sparse tree cover promotes the formation of a continuous grass layer (Archibald et al. 2009; Hennenberg et al. 2006; Pueyo et al. 2010), in turn limiting the density of trees in the landscape (Higgins et al. 2000; Prior et al. 2010; Staver et al. 2009). Meanwhile, dense tree cover shades out grasses, resulting in a discontinuous grass layer that impedes fire spread (Archibald et al. 2009; Hennenberg et al. 2006; Pueyo et al. 2010). Here, we capture these two alternative feedbacks using a step fire-mortality function ' (see also Staver et al. 2011b; Staver and Levin 2012), that takes a high value (combining high fire frequency with its potential effects on forest trees) at low tree-cover and a low value (representing a background mortality rate in the absence of fires) at high tree-cover. Finally, we assume that in the absence of fire, the tree cover accumulates logistically to some carrying capacity, with a per-capita growth rate that we normalize, without loss of generality, to precipitation *P*, reflecting an increase in tree growth rates [via increased primary productivity (Lieth 1975)] with increasing precipitation.

With these simplifying assumptions, the mean-field or the reaction term can be mathematically expressed as

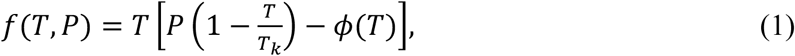

where *T, T_k_* and *P* represent tree cover, local carrying capacity, and precipitation, respectively. This reaction term has two important ecological features. First, in the absence of fire (*i.e.*, at *φ* = 0), the system equilibrates to a high tree-cover state. This feature of the reaction term *f*(*T, P*) is consistent with long-term (50-60 years) fire experiments that show that active fire suppression in mesic savannas can result in a closed canopy forest (Bond et al. 2005; Swaine et al. 1992; Trapnell 1959). Second, in the presence of fire, the mortality rate of trees has a threshold response to the tree cover itself (fire-vegetation feedbacks), consistent with previous empirical work (Archibald et al. 2009), because of which the equilibrium tree cover becomes bimodal in some intermediate range of rainfall. Theoretically, this implies that inclusion of fire results in a potential decrease in tree cover below the system’s carrying capacity by allowing for multiple stable states, corresponding to savanna 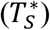 and forest 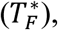 for some parts of parameter space. This is also evident from the bifurcation diagram in figure 1, which shows that both savanna and forest are stable states in the intermediate precipitation region, bounded by the two critical precipitation values (*P_SF_* and *P_FS_*); meanwhile, outside this rainfall region, the system has only one stable solution corresponding to savanna and forest in low and high precipitation regions, respectively. An analogous mean-field system has been thoroughly elaborated in a number of papers (Staver et al. 2011b; Staver and Levin 2012; Touboul et al. 2018).

**Figure 1:**
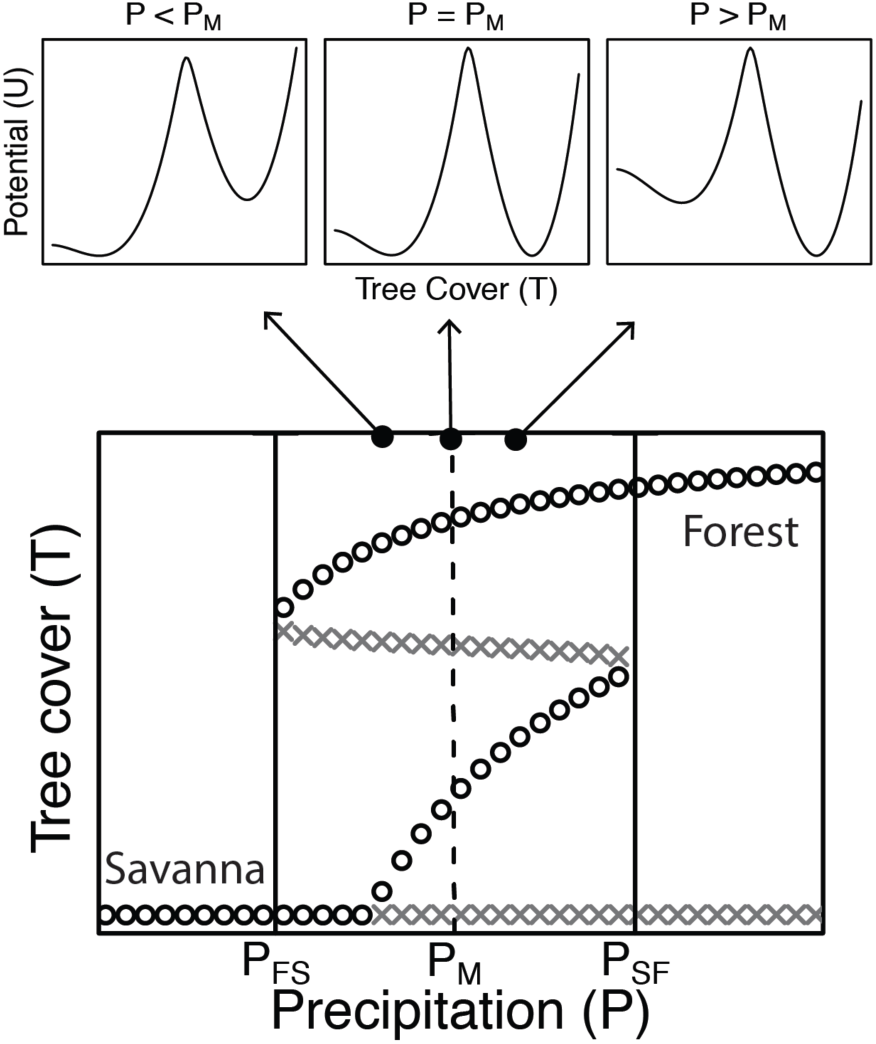
Potential functions of different vegetation configurations (top) and bifurcation diagram of the mean-field reaction term (bottom). In the intermediate precipitation region (bounded by the critical precipitation values *P*_FS_ and *P*_SF_), the bifurcation diagram shows that the system can exist in both savanna and forest states, depending upon the initial conditions. In the bistable region, the depth of the potential function (U in the top row) corresponding to savanna and forest states depends on the precipitation value. Both savanna and forest states have equal potential at a unique precipitation value, referred to as Maxwell precipitation (*P_M_*). Below (above) *P_M_*, savanna (forest) state has a deeper potential than forest. In the bottom panel, stable (unstable) equilibrium points are marked as black circles (dark-grey crosses).

Next, we incorporate seed dispersal in our model following Skellam (1951). In his paper, Skellam (1951) assumed that a plant disperses its propagules like a random walk process, with the probability of finding a propagule highest near the parent stem and falling off with increasing distance (Levin et al. 2003; Okubo and Levin 2013). This movement of plant populations, although random at an organism level can be statistically approximated to a continuous Diffusion (or Laplacian) operator ∇^2^ when scaled to the landscape level (Okubo and Levin 2013; Skellam 1951). Mathematically, ∇^2^ is defined as 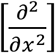 in 1D and 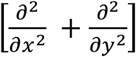 in 2D. Here, for simplicity, we assume that seed dispersal is isotropic, and ignore advective effects, for example due to wind.

Finally, combining the reaction (mean-field) and the diffusion (spatial) components of the model yields a reaction-diffusion equation:

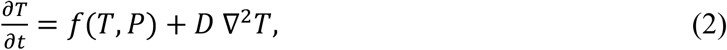

where *D* is the diffusion coefficient that captures the rate of seed spread.

Although the reaction-diffusion approach to model plant dynamics has proven to be quite useful because of its analytical tractability and mathematical simplicity, this approach has some inherent drawbacks, such as approximating discrete variables (such as habitats) as continuous (Keitt et al. 2001) and failing to consider the effects of long-range seed dispersal (Kot et al. 1996; see Appendix C), both of which have been previously shown to yield qualitatively different results. Moreover, we also ignore fire spread as spatially explicit process [see Schertzer et al. (2015) for a more realistic way of modeling fire spread within savannas and (Cochrane 2003; Cochrane et al. 1999) for discussion of the spatial structure of fire spread at the savanna-forest boundary]; instead, we incorporate fire effects only in the reaction term. Nevertheless, at continental scales, in the absence of appropriate continuum models, a diffusion model is a reasonable place to start.

In the next section, we explore the behavior of equation (2) using a series of simplifying assumptions that are ecologically relevant. It may also be worth mentioning that the qualitative behavior of the equation (2) is independent of the particular details of *f*(*T, P*). However, in this paper, we use a particular functional form of *f*(*T, P*) motivated by previous work on the subject (Staver et al. 2011b; Staver and Levin 2012). We do this to compare and contrast the simulation results of the previous mean-field model and its spatial counterpart, presented here.

## Methods and Results

Since the mathematical literature on bistable reaction-diffusion models is scattered across various subfields of physics (Coleman 1977), mathematics (Aronson and Weinberger 1975; Bramson 1983; Fife and McLeod 1977), and ecology (Lewis and Kareiva 1993; Murray 2001; Okubo and Levin 2013), we begin by summarizing some of the well-known results of the 1D diffusion model in the context of savanna and forest biomes. Although some results for the 1D model have been presented numerically elsewhere (Eby et al. 2017; van de Leemput et al. 2015), here we provide analytical results that may yield deeper insights. These results will also provide a baseline for comparison with the 2D diffusion model that has not been discussed in the literature.

### Reaction-Diffusion Model in One Dimension

Since one of the primary goals of the paper is to determine the spatial limits (boundaries) of savanna and forest biomes, it is natural to look for solutions that naturally give rise to boundaries. Based on the extensive literature on invasion biology (Hastings et al. 2005; Keitt et al. 2001), we know that equation (2) has a traveling wave solution (see Fig. A1), where the wavefront can be interpreted as the savanna-forest boundary. In this section, we find the velocity of movement of the savanna-forest boundary as a function of system parameters, e.g., precipitation. We then set the velocity to zero to find the equilibrium boundary position.

Using the generalized waveform in one-dimension, *T*(*x, t*) = *T*(*x − vt*) (Aronson and Weinberger 1975; Bramson 1983; Fife and McLeod 1977; Murray 2001; Okubo and Levin 2013), where D is the velocity of the savanna-forest boundary, we find that

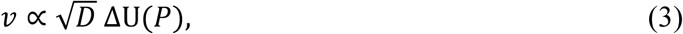

where ΔU(*P*) is defined as the difference between values of the potential function at forest and savanna state, respectively (see Appendix B.1 for calculations). Mathematically, the potential function is defined as 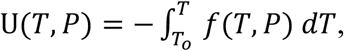 where *f*(*T, P*) is the mean-field growth function (Nolting and Abbott 2016; Strogatz 2014). This potential function is a formal way of defining the concept of a potential landscape that is commonly used to understand the resilience of dynamical systems (Holling 1996; Strogatz 2014). In a bistable system, the potential landscape consists of two wells corresponding to the two stable states of the system (see top row in Fig. 1). In the equation above 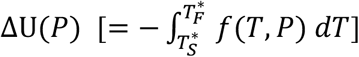 is the difference between the depth of potential wells corresponding to savanna and forest.

The equation above suggests that the magnitude of D is proportional to diffusion 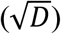 and the difference in the depth of the potential wells (ΔU), while the direction of *v* is purely determined by the sign of ΔU(*P*). Thus, in a homogeneous landscape (*e.g.*, with constant precipitation across the whole landscape), the state with lower potential invades the one with higher potential, except in the trivial case when the potential for both states is equal (*i.e.,* ΔU = 0; see top row in Fig. 1). The trivial case occurs at a unique precipitation value, which is referred to as Maxwell precipitation (*P*_M_) (Bel et al. 2012; Boettiger and Hastings 2013; Carr et al. 1984; Clerk-Maxwell 1875; Martín et al. 2015; Pomeau 1986; van de Leemput et al. 2015; Weissmann and Shnerb 2014; Wuyts et al. 2017; Zelnik and Meron 2018). Next, to obtain the velocity of movement of the savanna-forest boundary as a function of precipitation, we Taylor expand ΔU(*P*) in equation (3) around +_N_:

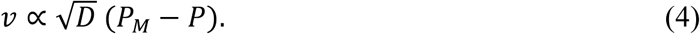

This equation implies that in a landscape with precipitation greater than *P_M_*, forest encroaches savanna (*v* < 0), and conversely, that in a landscape with precipitation less than *P_M_*, savanna encroaches forest (*v* > 0). Only when the landscape receives precipitation exactly equal to +_N_ is the savanna-forest boundary neutrally stable (*i.e.*, the magnitude of the small perturbations to the boundary neither increases or decreases over time). In other words, in a homogeneous precipitation landscape with *P* ≠ *P_M_*, a stable savanna-forest boundary is not possible, under these assumptions (van de Leemput et al. 2015).

### A Precipitation Gradient and Stable Savanna-Forest Boundary

In the previous section, we assumed homogeneous precipitation conditions. However, at continental scales, landscapes have precipitation gradients. In this section, we show how a precipitation gradient can lead to a stable savanna-forest boundary (van de Leemput et al. 2015; Wuyts et al. 2018).

To do this, we consider a 1D landscape with a linear precipitation gradient with precipitation *P* at site *x* given by

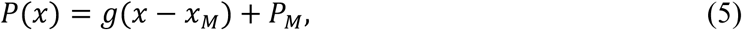

where *x_M_* is the spatial location receiving +_N_ and S is the change in precipitation per unit distance (precipitation gradient constant). Substituting equation (5) into equation (4) we get

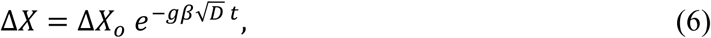

where *X* is the position of the savanna-forest boundary, Δ*X* = *X* − *x_M_* is the deviation of the boundary from *x_M_*, and *β* the natural logarithm of the proportionality constant in equation (4).

Equation (6) highlights two important features of the savanna-forest boundary. First, in a 1D landscape with linear precipitation gradient, the boundary equilibrates to *P_M_* (Fig. 2). Second, if the boundary is perturbed locally (in any direction), it will recover back to *x_M_*. Moreover, the characteristic timescale of recovery is inversely proportional to 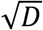 precipitation gradient constant *g* . This suggests that the savanna-forest boundary is resilient to local spatial perturbations (Fig. 2). Although not shown here, our numerical experiments in 1D also suggest that the equilibrium distribution of savanna and forest is independent of initial conditions (see also van de Leemput et al. 2015; Wuyts et al. 2018).

**Figure 2:**
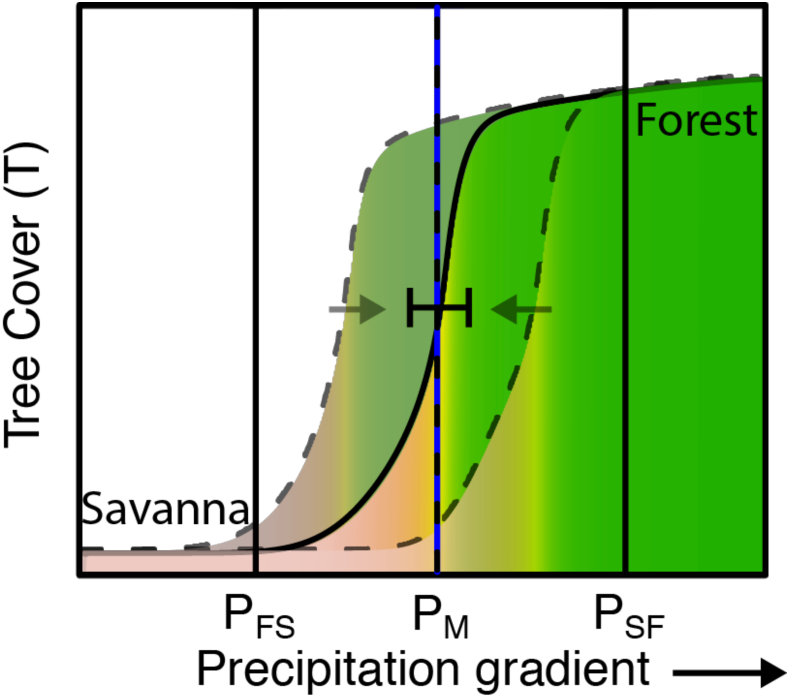
Equilibrium (solid sigmoidal curve) and transient (dash sigmoidal curve) tree cover along a linear spatial precipitation gradient in a 1D landscape. The two solid vertical lines correspond to the two critical points (*P*_FS_ and *P*_SF_), and the vertical black dashed line to the Maxwell precipitation (*P_M_*). The plot suggests that spatial interactions coupled with a large-scale gradient in precipitation can result in the spatial aggregation of savanna and forest, separated by a stable savanna-forest boundary (indicated by the blue vertical line, in this case coincident with *P_M_*). This boundary is resilient to perturbations and always recovers back to its equilibrium position after a disturbance. This model, however, fails to reproduce the non-deterministic relationship between biome and precipitation, observed in the empirical data. The simulations were initialized with random initial condition (see Numerical Methods in Online Appendix D).

Thus, the 1D diffusion model reproduces what the bistable biome theory (Beckage et al. 2009; Staver et al. 2011b; Staver and Levin 2012) could not: that spatial interactions, overlaid on a large-scale precipitation gradient, can result in the spatial aggregation of savanna with savanna and forest with forest, separated by a stable savanna-forest boundary (Fig. 2). Moreover, the model also predicts that biome shifts are reversible provided the climatic conditions are restored. Unfortunately, the 1D diffusion model also predicts that the spatial limits of savanna and forest biomes are solely determined by Maxwell precipitation (*P_M_*). In other words, this model fails to produce overlap in the rainfall ranges of savanna and forest biomes (see also Eby et al. 2017; van de Leemput et al. 2015; Wuyts et al. 2018), observed in the empirical data (Dantas et al. 2016; Hirota et al. 2011; Staver et al. 2011a; Staver et al. 2011b).

### Reaction-Diffusion in Two Dimensions

Above, we assumed a one-dimensional landscape. This assumption, however, may not be realistic for understanding distribution of savanna and forest biomes, since it is somewhat obvious to observe that their dynamics are better described on a two-dimensional landscape. In this section, we show that adding a second dimension can qualitatively change the equilibrium position of the savanna-forest boundary, which can explain the overlap in the rainfall ranges over which biomes occur.

To incorporate the second dimension in the model, we use a 2D polar representation of the Laplacian operator in equation (2). Following the same analytical approach as in the 1D case, we show that

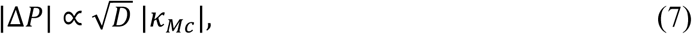

where Δ*P* represents the difference between the precipitation at the savanna-forest boundary and the Maxwell precipitation contour (*P_MC_* notationally distinct from the Maxwell point in 1D *P_M_* see Appendix B.2 for calculations), |*κ_MC_*| the absolute curvature of *P_MC_* (a measure of roundness; see Fig. A2), and A the diffusion constant (as above). This equation describes the local deviation of the savanna-forest boundary from the location of the *P_MC_* in terms of the difference in the precipitation; to obtain the deviation in terms of absolute distance, we multiply Δ*P* by *g* (precipitation gradient; see Eq. 5).

In plain terms, this means that, in a 2D landscape, the location of the boundary between savanna and forest is not determined only by precipitation but also crucially depends on the geometrical shape of the precipitation contours (specifically, of the Maxwell precipitation contour *P_MC_*). When the Maxwell precipitation contour is a straight line (*i.e.*, where |κ*_MC_*| ≠ 0), the system behaves like a 1D model, and the savanna-forest boundary coincides with *P_MC_* (Fig. 3A). However, for an arbitrarily shaped *P_MC_* (|κ*_MC_*| ≠ 0), the boundary deviates from the *P_MC_* depending upon the local curvature of *P_MC_* (Fig. 3B).

**Figure 3:**
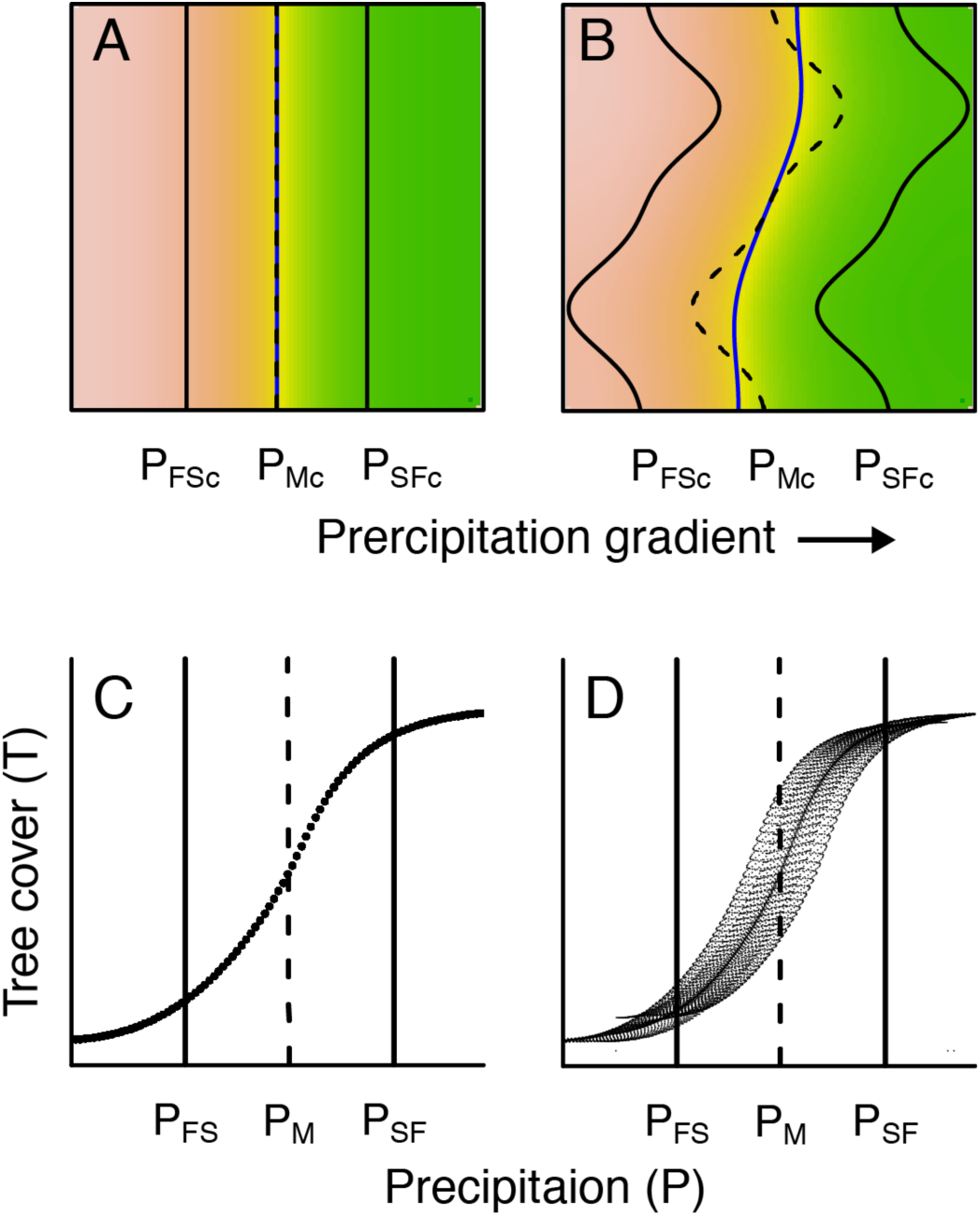
Simulated equilibrium distribution of tree cover (A and B) and its relationship with precipitation (C and D) in a 2D landscape witha monotonic precipitation gradient. The columns show the results for two geometries of precipitation contours: linear (A and C) and curved (B and D). Simulations suggest that in a 2D landscape, the equilibrium position of the boundary is not only determined by *P_M_*, but also depends on the curvature of the Maxwell precipitation contour κ_MC_ . When κ_MC_ = 0 (linear *P_MC_*), the boundary aligns with *P_MC_* (A). This situation is analogous to the one-dimensional model in figure 2. However, when κ_MC_ ≠ 0 (arbitrary shaped *P_MC_*), the boundary deviates from *P_MC_* according to equation (7) (B). These curvature effects can reproduce the non-deterministic relationship between biome and precipitation (A), missing in the one-dimensional model (Fig. 2). The simulations were initialized with random initial condition (see Numerical Methods in Online Appendix D).

Ecologically, curvature effects described in equation (7) arise because of source-sink dynamics (Pulliam 1988; Pulliam 2000) at the savanna-forest boundary. When *P_MC_* is a straight line, the inflow and outflow of seeds are balanced, thus resulting in a stable savanna-forest boundary that coincides exactly with *P_MC_*. However, if *P_MC_* is curved, the balance between inflow and outflow of seeds is disrupted. For example, when the shape of *P_MC_* is such that there are more forest neighbors than savanna neighbors surrounding a point in the landscape with *P* = *P_MC_* (upper part of Fig. 3B), the inflow of seeds will be higher than their outflow. This creates a net positive inflow of seeds, resulting in a higher growth rate of trees. Forests expand, pushing the boundary into savanna region till the added growth rate of trees due to a higher influx of seeds is compensated by reduced growth rate due to a decrease in precipitation. Conversely, when a point on *P_MC_* is surrounded by more savanna patches than forest patches (lower part of Fig. 3B), there will be a net positive outflow of seeds, which will favor savanna expansion. Similar to the previous case, the boundary will move into forest regions until the reduced growth rate of trees due to a lower influx of seeds is balanced by increased growth rate due to an increase in precipitation.

The reaction-diffusion model, presented above, however, has some assumptions that are likely to be violated in real-world: (1) that dispersal is local (because of diffusion approximation), (2) that vegetation dynamics have no demographic or external noise, and (3) that the reaction part has a well defined potential function. As a robustness check, we relax these assumptions one by one, and numerically test their consequences for the theoretical results presented above. First, we find that incorporating long-range dispersal (via fat tail dispersal kernels) does not change equilibrium biome distributions (see Appendix C), presumably because fire vegetation feedbacks prevent tree establishment far away from the source even when a seed arrives there (Barton and Turelli 2011; Bates et al. 1997; Kot et al. 1996). Second, we find that adding noise makes the boundary increasingly rough with increasing noise; however, the location of the boundary at a coarser scale does not move appreciably from its equilibrium position predicted from the deterministic 2D diffusion model (see Fig. D1). Thirdly, and finally, we consider a two-dynamical-variable reaction-diffusion system where a potential function cannot be defined. In such a system, the position of the boundary, in addition to the control parameter (*e.g.*, precipitation), is dependent on the ratio of the two diffusion constants (Fig. D2); the system still exhibits curvature effects in equation (7). Therefore, in no case did we find that violating the above assumptions qualitatively changed dynamics.

To summarize, curvature effects in the 2D diffusion model can phenomenologically reproduce the overlap in the precipitation ranges over which savanna and forest biomes occur, missing from the 1D diffusion model, while simultaneously retaining the spatial aggregation property of biomes (Fig. 3D). Moreover, the 2D diffusion model suggests that this precipitation overlap is not maintained by hysteresis, a defining feature of a bistable biome theory. Instead, our simulations and analytical calculations suggest that in a landscape with a monotonic precipitation gradient, hysteresis is unlikely. Below we discuss an ecological scenario under which hysteresis may reappear.

### Critical Patch Size Effects in a Landscape with Non-Monotonic Precipitation Gradient

In the previous section, we assumed a monotonic gradient in precipitation. However, precipitation gradients in real-world landscapes are not always monotonic. As such, a landscape can have a complex distribution of precipitation with high precipitation regions intermittently distributed in low precipitation regions, and vice versa. In the following section, we describe how this feature of precipitation gradients can potentially lead to hysteresis.

But before we do that, we first consider a simpler case of a homogeneous precipitation model for analytical insight. Based on the theoretical works of Bradford and Philip (1970a,b), it can be shown that in a homogeneous precipitation landscape with bistable dynamics, the fate of an invasion process by a particular vegetation state into another is dependent on two factors: precipitation + and the initial patch area of the invading state _. An invading patch of vegetation smaller than a critical patch area (*A* < *A_c_*) will not be able to expand even though that vegetation state is climatically favourable, *i.e.,* the state which has lower potential (Bradford and Philip 1970a; Bradford and Philip 1970b; Holmes et al. 1994; Oxtoby 1998; Skellam 1951). Our calculations suggest that near +_N_, the critical patch size in a homogeneous landscape can be approximated as

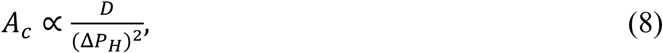

where Δ*P_H_*is the difference between the precipitation of the homogeneous landscape and *P_M_*. To fully understand the implications of equation (8), consider an initial savanna landscape with precipitation just above *P_M_*. Although in this landscape forest is more favourable than savanna because of lower potential (Eq. 3), the forest state will only be able to invade if there is an initial patch of forest that has an area greater than *A_c_* (Eq. 8). This is because a small patch of forest has a high perimeter-to-area ratio (Skellam 1951), such that the accumulation of trees is slow because seed inflow per unit area from forest patches is low, preventing forest expansion. By the same token, a landscape in a forest state with precipitation just below *P_M_* would require a large patch of savanna to overcome high levels of seed rain from neighboring forest patches.

The same phenomenon also applies to a landscape with precipitation gradients. Consider an initial savanna landscape with spatially varying precipitation patterns such that the whole landscape has rainfall less than *P_M_*, except in the center where the rainfall is just above *P_M_*. Since the whole landscape was initialized with savanna, the center of the landscape will remain in a savanna state, unless the central region is initialized with forest patch of area greater than *A_c_*. Conversely, a similar argument holds for an all forest landscape with a low-rainfall island in the center.

This suggests that the vegetation state of small and isolated patches in intermediate rainfall regions depends on the availability of a nucleation center, suggesting that the characteristic biome state in those areas might be contingent on historical biome patterns, thus exhibiting hysteresis. And more importantly, this analysis suggests that critical patch size (Eq. 8), in addition to curvature effects (Eq. 7), can also explain overlap in the rainfall ranges of savanna and forest biomes.

#### Curvature and Critical Patch Size Effects in Empirical Systems

In this section, we test some of the key analytical predictions of our 2D reaction-diffusion model—particularly those concerning curvature (Eq. 7) and critical patch size effects (Eq.8)—using real savanna and forest distributions in Sub-Saharan Africa. As described above, a 2D reaction-diffusion approximation of savanna-forest dynamics predicts that the location of the savanna-forest boundary with respect to precipitation should vary depending upon the local curvature of the boundary (Eq. 7; Fig. 3B) in such a way that the difference between the precipitation at the boundary and *P_MC_* is linearly proportional to the local curvature of the boundary. As such, plotting the absolute curvature of the boundary (ignoring its convexity) as a function of boundary precipitation should yield a V-shaped curve, the vertex of which should correspond to *P_M_*. Indeed, using African tree cover and mean annual precipitation (MAP) data (see Data Analysis in Online Appendix D), we show that the current distribution of savanna and forest in Africa is consistent with this prediction (black line in Fig. 4). The location of the vertex of that curve provides an estimate of *P_M_* = 1508 ± 84 mm MAP, consistent with previous ^empirical work (Staal et al. 2016 found *P_M_* = 1580 mm MAP). The confidence interval for this^ estimate was determined by calculating *P_M_* for various combination of parameter values of (*i*) boundary tree cover (73-80%) used for identifying boundary location, and (*ii*) arc length of the boundary (100-1000 km) used to estimate curvature. For more details on estimating *P_M_*, see figure D3 and sensitivity analysis in Online Appendix D.

**Figure 4:**
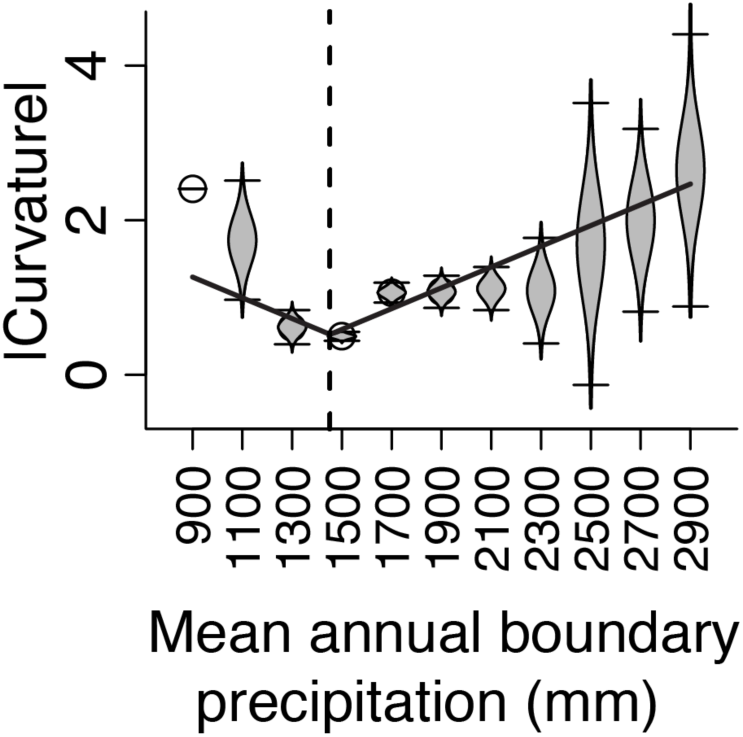
Empirical response of local savanna-forest boundary curvature to mean annual precipitation at the boundary in sub-Saharan Africa. Results show that absolute curvature vs. boundary precipitation exhibit V-shaped relationship (black line), consistent with our theoretical prediction (eq. 7). In theory, the vertex of V corresponds to *P_M_*, loosely corresponding to results from the extensive sensitivity analysis that estimates *P_M_* = 1508 ± 84 MAP (Fig. D3; see Data Analysis in Online Appendix D).

Next, we compare the above estimate of *P_M_*, by estimating *P_M_* with an alternative method. This involves simulating the potential distribution of savanna and forest using a 2D reaction-diffusion model with current biome distributions as the initial condition. To do this, we simulated spatial distributions of savanna and forest using present-day precipitation patterns for various combinations of *P_M_* and A (see Data Analysis in Online Appendix D), and, using a genetic algorithm (Scrucca 2013), selected those parameter values (*P_M_* and *D*) that yielded the ‘best fit’ to the current distribution of biomes. Here, we refer to ‘best fit’ as maximizing pixel by pixel match between simulated and empirical savanna and forest distribution (see Data Analysis in Online Appendix D). This procedure yielded an estimate of *P_M_* (= 1538 mm MAP) that lay within the expected precipitation range obtained from the curvature analysis in figure 4 (see Fig. D4). These large-scale simulations also reproduced empirically observed biome distributions in Sub-Saharan Africa surprisingly well for such a simple model (Fig. 5A), except for small regions in the Bateke Plateau in Congo and Western Africa. In the Bateke Plateau, empirically observed savannas may be maintained because shallow sandy soils that reduce effective soil moisture (White 1986) or may alternatively be anthropogenic. Meanwhile, it is well established that savannas in Western Africa are a result of historical deforestation (Adams and Faure 1997; Aleman et al. 2017).

**Figure 5:**
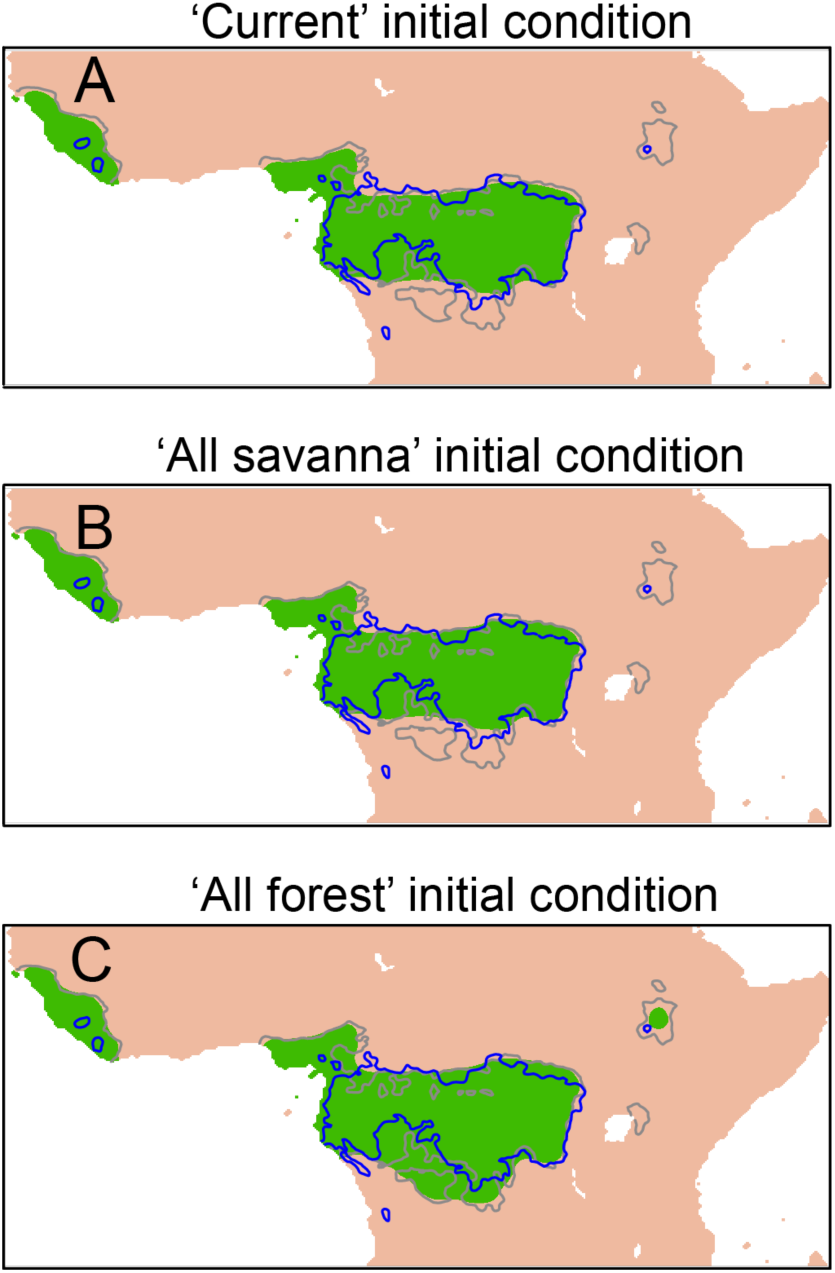
Simulated distributions of savanna and forest in Africa, initialized with the current distribution of biomes (A), all savanna (B), and all forest (C). Blue lines correspond to the observed present-day savanna-forest boundary, and the grey line represents *P_MC_* (with *P_M_* = 1538 mm MAP). The large-scale simulations in (A) and (B) matched, and reproduced the current distribution of savanna and forest, except the edaphic savannas on the Bateke Plateau in Congo and deforested areas in western Africa. However, the simulations in(C) significantly overpredicted the extent of forest in the Southern Congo. This region can climatically support both savanna and forest depending upon the historical vegetation state of the region (see Data Analysis in Online Appendix D).

Next, to check whether the results of large-scale simulation in figure 5A were dependent on the initial conditions – at least theoretically possible because of critical patch size effects, as described above – we simulated the vegetation distribution for two more initial conditions: ‘all savanna’ and ‘all forest’ in Sub-Saharan Africa (Fig. 5B-C), using the best fit parameter values estimated above (*P_M_* and *D*). Simulations with ‘all savanna’ initial conditions (Fig. 5B) matched those using current distributions as initial conditions (see Fig. 5A). However, ‘all forest’ initial conditions produce substantially different biome patterns in the Southern Congo and Ethiopian Highlands (Fig. 5C).

We propose that the critical patch area (*A_c_*) requirement can potentially explain why the simulations over-predict the forest extent in the Southern Congo (Fig. 5C) and under-predict the forest extent in Ethiopian Highlands (Fig. 5B). Since both of these regions are disconnected from the main forest cluster by savanna vegetation, biome distributions in these regions are dependent on the availability of historical nucleation centers (or initial conditions; Eq. 8). Based on our simulations we suspect that Southern Congo and Ethiopian Highlands were historically occupied by savannas and forests, respectively, which resulted in their present distribution. Although this claim is currently hard to test due to lack of reliable long-term paleo-records from these regions (however, see Elenga et al. 1994; Jolly et al. 1998), historical vegetation reconstructions for the early 20^th^ century (Aleman et al. 2017; White 1986) are consistent with the theoretical predictions of the model.

### Comparisons with Alternative Models

Our calculations show that novel dynamical features of the 2D diffusion model — such as spatial aggregation (Eq. 6), curvature effects (Eq. 7), and critical patch size (Eq. 8) — can qualitatively explain many empirical features of savanna and forest distributions that previous biome distribution models could not. In this section, we investigate whether these dynamical features improve upon the predictions from previously proposed models of biome distribution. To do this, we simulate the distribution of savanna and forest in Sub-Saharan Africa using three alternatives (see Data Analysis in Online Appendix D). First, (*a*) we consider a ‘one-climate one-biome’ model in which the savanna-forest boundary is determined by a unique precipitation contour. This model is analogous to the classical biome theory (Fig. 6A). Next, (*b*) we consider a model in which the local vegetation dynamics in each patch are governed by mean-field bistable model and the neighbouring patches do not interact. In this model, we randomly initialize the landscape with savanna and forest patches (Fig. 6B); note, however, that this test does not consider the possibility that initial conditions could be spatially structured, leading to spatial structure in biome distributions today. Finally, (*c*) we consider a 2D reaction-diffusion model, already described at length above (Fig. 6C). The diffusion model incorporates both bistability vegetation dynamcis and dispersal.

**Figure 6:**
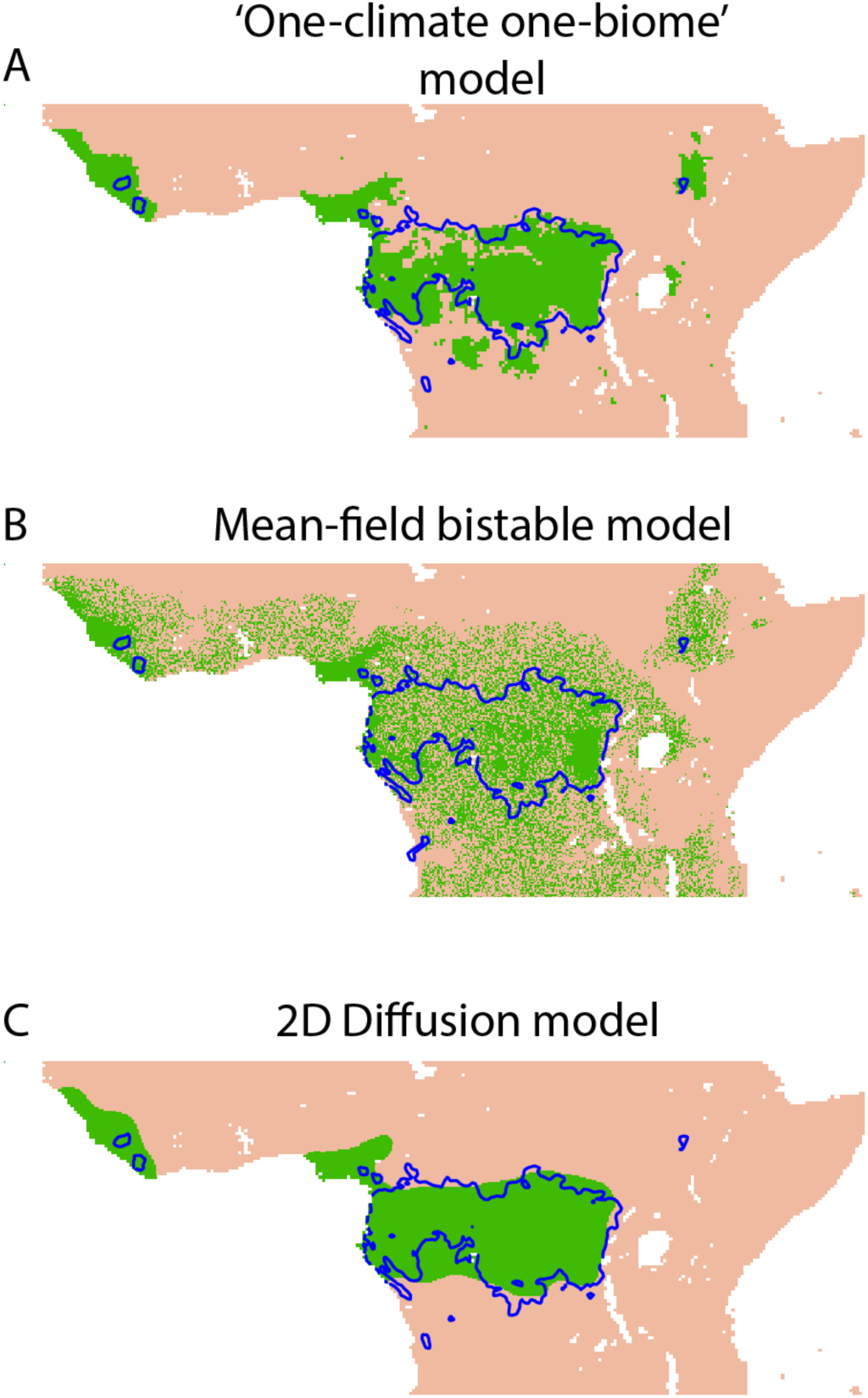
Simulated distribution of savanna and forest biomes from three biome distribution models: ‘one-climate one-biome’ model (A), mean-field bistable model (B), 2D reaction-diffusion model (C). Blue lines correspond to the present-day savanna-forest boundary. These results indicate that the 2D reaction-diffusion model can, with tuning, describe the quantitative distribution of biome patterns. This model reproduces both spatial aggregation and overlap in rainfall ranges of biomes and is also the best predictor of biome patterns in Central Africa (see Table 1 and Data Analysis in Online Appendix D).

We measure whether these models can – with parameter optimization – reproduce three components of biome distribution: overlap in the rainfall ranges of biomes, the spatial aggregation of savanna with savanna and forest with forest, and the match between the simulated and actual distribution of biomes (see Fig. 6, Fig. D5, and Table 1). Note, again, that tuned parameters do not necessarily correspond to demographic rates, etc., that might be measured empirically; note also that the three model alternatives we propose here are not exhaustive.

**Table 1:**
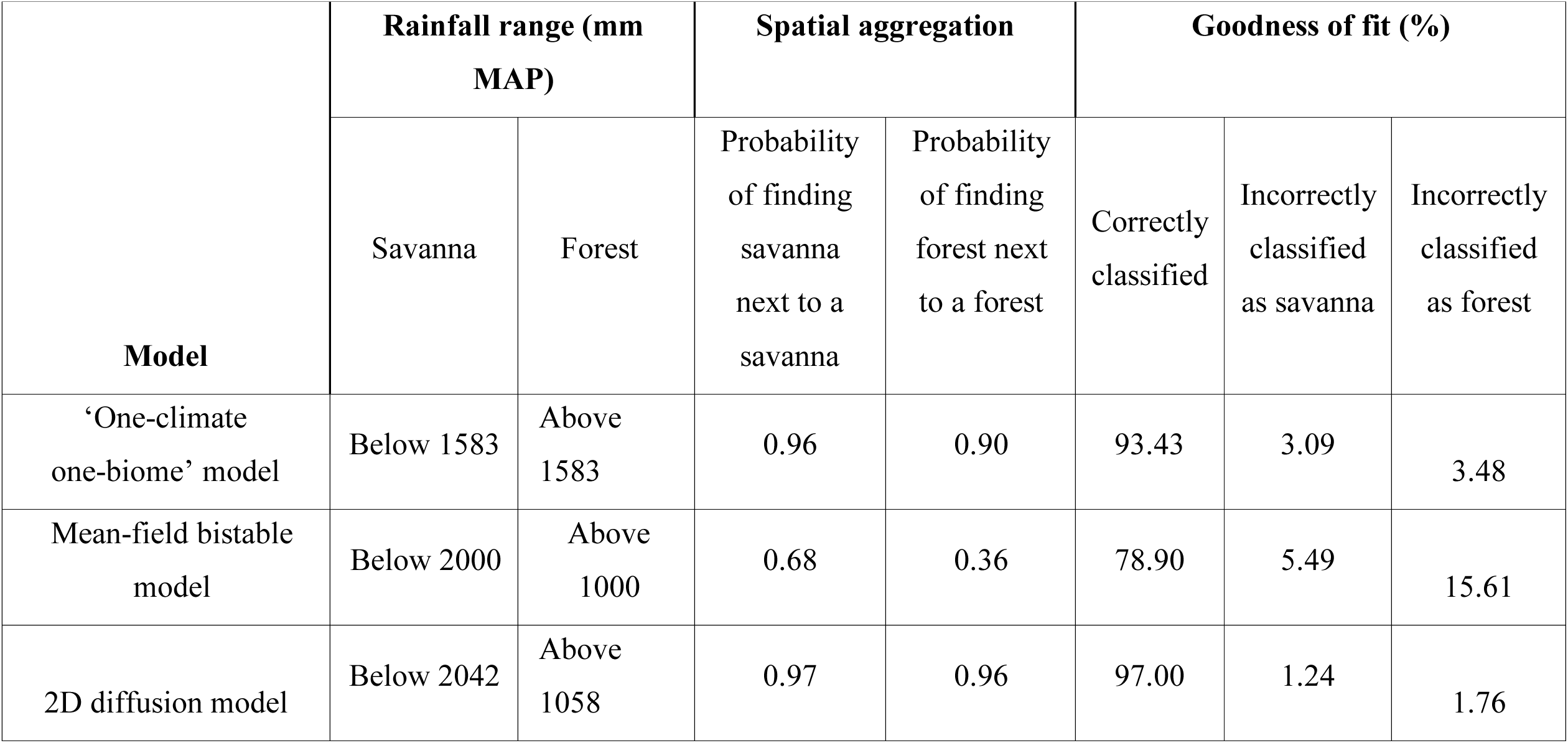
Summary statistics of the simulated distribution of biomes in Central Africa using three alternative models. The performance of models was evaluated on three aspects: overlap in the rainfall ranges of biomes (columns 2 and 3), spatial aggregation of savanna with savanna and forest with forest (columns 4 and 5), and match between the simulated and actual distribution of biomes (columns 6, 7, and 8). Note that we excluded the contributions of the deforested regions in Western Africa and edaphic savannas of Bateke Plateau while calculating the goodness of fit (see Data Analysis in Online Appendix D).

In the ‘one-climate one-biome’ model, the precipitation cutoff between savanna and forest was found to be 1583 mm MAP (see Table 1). Whereas, in the other two models the rainfall ranges of savanna and forest showed considerable overlap between 1000 mm and 2000 mm. Meanwhile, the one-climate one-biome and 2D diffusion models show a high probability of spatial aggregation (above 90%) that is missing in the mean-field bistable model (below 68%).

Thus, of the three models, only the 2D diffusion model can reproduce both spatial aggregation and overlap in the rainfall ranges of biomes. Therefore, it is not surprising that the 2D diffusion model also outperforms (97% accuracy) other models in terms of predicting the spatial distribution of biomes in Sub-Saharan Africa.

In summary, all models, except in a one-climate one-biome model, reproduce at least some overlap in the rainfall ranges of biomes (Table 1 and Fig. D5). Of the two remaining models, the mean-field bistable model fails to reproduce the spatial aggregation of biomes (Fig. 6 and Table 1). This leaves us with the 2D reaction-diffusion model, which reproduces not only the climatic overlap in the limits of biomes, and the spatial aggregation in biome distributions, but also the overall biome distributions in Sub-Saharan Africa with remarkable accuracy (see Table 1).

## Discussion

In this paper, we develop and analyze a reaction-diffusion model to examine the contributions of dispersal to the distribution and resilience of tropical savanna and forest biomes. The model assumes that the local mean-field dynamics of biomes are governed by non-linear fire-vegetation feedbacks and that adjacent savanna and forest patches interact spatially through seed dispersal.

We find that the model reproduces empirical features missing from existing biome distribution models. Specifically, the 2D reaction-diffusion model simultaneously reproduces both overlap in the climatic ranges of biomes, as well as spatial aggregation of savanna with savanna and forest with forest. As before, we find that fire-vegetation feedbacks may substantially expand savanna distributions at the expense of forests, but that in a spatial context this does not necessarily translate into bistable vegetation distributions. Instead, the equilibrium position of the savanna-forest boundary is determined by a combination of three factors: (a) climate (via impacts on the relative depth of potential wells for each biome), (b) source-sink dynamics (via local curvature of the Maxwell precipitation contour), and, occasionally, by (c) availability of historical nucleation centers (which contributes an element of hysteresis to distribution dynamics, albeit more limited than that described before). These theoretical predictions are empirically consistent with observations of the curvature of the savanna-forest boundary, and large-scale simulations which show that the 2D diffusion model can — with parameter optimization—reproduce empirically observed patterns of savanna and forest distributions in Sub-Saharan Africa.

These findings have direct implications for how we think of the stability and resilience of tropical biomes. Classical biome theories suggest that perturbations to biome distributions should be easily reversible since vegetation tracks climate directly (Holdridge 1947; Schimper 1902; von Humboldt and Bonpland 1807; Whittaker 1970). By contrast, more recent work has suggested that fire-vegetation feedbacks can stabilize savanna as an alternative to forest in some areas, such that perturbations to biome distributions may not be easily reversible (Beckage et al. 2009; Staver et al. 2011b; Staver and Levin 2012). Here, we show that combining a spatial dispersal process with an underlying bistable model radically alters stability predictions: biome recovery after perturbation becomes much more likely, even if fire-vegetation feedbacks do modify vegetation (which they probably do; see Bond et al. 2005). In this scenario, biome transitions may be regionally predictable and reversible, even if they are locally abrupt.

However, there is a notable caveat to this prediction. In isolated rainfall islands, vegetation distributions may exhibit hysteresis; analogously, if remnant vegetation patches are reduced below a critical area, recovery of the boundary may be impossible, resulting in a permanent loss of vegetation. As a result, extensive historical forest loss, *e.g.*, in West Africa, coastal Kenya and Tanzania, and the Ethiopian highlands, may be irrecoverable without direct intervention, since remnant forest patches may be too small for forest to recolonize successfully (Aleman and Staver 2018). This also raises contrasting concerns about proposed afforestation plans in mesic savannas of the Southern Congo (Veldman et al. 2015). These isolated mesic savannas might be historically maintained as a stable alternative biome state (Aleman et al. 2017); proposed afforestation practices (Veldman et al. 2015) in these regions could trigger a permanent shift in ecosystem state from savanna to forest (Fig. 5C), which may lead to loss of endemic biodiversity in mesic savannas (Bond 2016) and wastage of scarce management resources.

The results of the reaction-diffusion model presented herein should be interpreted with caution, however. For starters, we here incorporate only a subset of important spatial processes, notably ignoring the long-range spread of fire within savannas (Schertzer et al. 2015) and local fire spread at the savanna-forest boundary (Cochrane 2003; Cochrane et al. 1999) both of which may be significant (note, however, that we have included fire effects in the reaction term of the model). However, ongoing analytical work on a more thorough set of models that examine fire effects at the boundary between savanna and forest (Durrett and Ma 2018) suggest that the phenomenological results presented herein may be applicable more broadly: scaling limits to those models appear also to be characterized by traveling waves, with the occurrence of stationary savanna-forest boundaries only in landscapes that include a gradient in rainfall (Durrett and Ma 2018). Notably, however, long-range fire effects seem to change predictions somewhat (Li et al., in review), resulting in the emergence of stable savanna-forest mosaics even under homogenous climatic conditions (Schertzer et al. 2015).

Another major question surrounds the problem of time-scales of ecological processes. Here, we have considered only the equilibrium distribution of biomes, ignoring the speed of equilibration. Modern climate change is sufficiently rapid (Karl and Trenberth 2003), and so associated with extreme climatic events (Jentsch et al. 2007; Katz and Brown 1992), that biome responses to ongoing anthropogenic global change are unlikely to be dominated by these local spatial processes. This may result in transient mismatches between climate and equilibrium vegetation (Webb 1986), which may be persistent from timescales ranging from decades to millennia depending on the speed of ecological dynamics (Hastings 2004; Hastings et al. 2018; Hastings and Higgins 1994). Therefore, understanding how fast biomes respond to changing climate (empirically from the paleo-records) and using this information to incorporate dispersal into existing non-spatial biosphere models (Bond and Keeley 2005; Bond et al. 2003; Moncrieff et al. 2014; Scheiter and Higgins 2009; Scheiter et al. 2013) will be critical to generating informative predictions for the effects of anthropogenic global change on biome distributions.

Projections show that rapidly changing climate (Lewis et al. 2011; Nepstad et al. 2004) and land-use change (Aleman et al. 2016; Cochrane and Laurance 2002) are expected to result in large-scale biome shifts, which may yield huge economic and ecological losses. Here, we argue that, except in a few cases, dispersal can, in general, increase the resilience of tropical savanna and forest biomes to natural and anthropogenic disturbances (see also van de Leemput et al. 2015). However, recovery from disturbance could be slow, due to slow dynamics of biomes and anthropogenic or natural dispersal barriers.

## Supporting information

## Author contributions

NG and ACS designed this model based on a concept from ACS. NG and VG implemented the diffusion and integro-differential equation models. NG performed simulations and data analyses. ACS and NG co-wrote the manuscript with feedback from VG and SAL. All the authors contributed ideas and discussions.

## Acknowledgements

Funding for this work was provided by a grant from the NSF to ACS (#DMS-1615531) and SAL (#DMS 1615585) and by Yale University. VG is supported by DBT-IISc partnership program. We thank Thierry Emonet for help with technical problems simulating the integro-differential equation model. We also thank members of the Staver lab, especially Julie Aleman and Madelon Case, for manuscript feedback.

