## Supplementary material for "Dispersal increases the resilience of tropical savanna and forest distributions"

**Appendix A: Supplementary Figures**

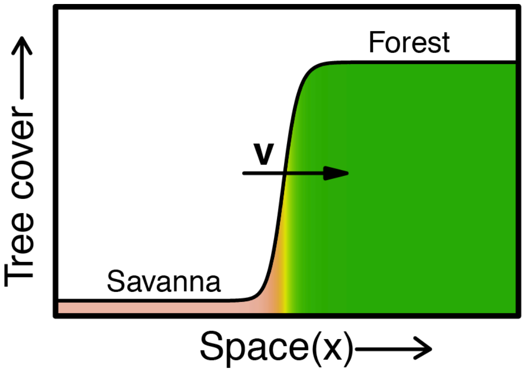


**Figure A1:** A schematic representation of a traveling wavefront. The wave of tree cover ($T$) is moving with a velocity $v$ in the positive $x$-direction. The left (right) part of the wave corresponds to low (high) tree cover savanna (forest). The vegetation wave is a monotonically increasing function of $x$, such that at the extreme ends of the landscape, $(T')^2 \rightarrow 0$, and at the boundary $(T')^2$ shows a sharp peak.

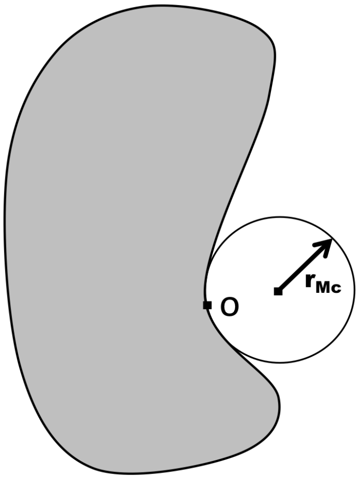


**Figure A2:** A schematic representation of radius of curvature ($r_{Mc}$) at a point $O$. Radius of curvature of a curve at a point is defined as the radius of the circle that passes through two arbitrarily close points on the curve. The curvature of a curve ($\kappa_{Mc}$) is mathematically defined as the inverse of $r_{Mc}$. Curvature of a straight boundary is 0.

### Appendix B: Supplementary Calculations

#### B.1 Speed of the savanna-forest boundary in a one-dimensional landscape

In the main text, we argued that equation (2) has a traveling wave solution. We refer the reader to the works of Aronson and Weinberger (1975); Bramson (1983); and Fife and McLeod (1977) for details of the proof and assume hereafter that equation (2) has a wave solution and can be represented as $T(x,t) = T(x - vt)$. Let $z = x - vt$, where $v$ is the velocity of the savanna-forest boundary. To avoid confusion, we will assume savanna encroaching forest as the positive direction as shown in figure A1. Mathematically, this implies that $T(z)$ is a monotonically increasing function such that $\lim_{z\to-\infty} T(z) = T_S^*$ and $\lim_{z\to\infty} T(z) = T_F^*$, where $T_S^*$ and $T_F^*$ are the stable roots of $f(T,P)$ corresponding to savanna and forest, respectively. Substituting $T(x - vt)$ in equation (2), we get

$$T'\,(-v) = f(T,P) + D\,T''. \qquad \text{(B1)}$$

Next, we multiply equation (1) by $T'$ and integrate it from $-\infty$ to $\infty$ w.r.t to $z$, to yield

$$-v\int_{-\infty}^{\infty}(T')^2 dz = \int_{-\infty}^{\infty} f(T,P)dT + [T']_{-\infty}^{\infty}. \qquad \text{(B2)}$$

The last term on the RHS goes to zero because at the extreme ends of the landscape the vegetation does not vary (Fig. A1), such that

$$v\int_{-\infty}^{\infty}(T')^2 dz = -\int_{-\infty}^{\infty} f(T,P)dT. \qquad \text{(B3)}$$

Next, we note that at the extreme ends of the landscape, the system is occupied by savanna and forest state (Fig. A1). In other words, we can replace the limits of the integral on the RHS with the density of savanna and forest states, to yield

$$v\int_{-\infty}^{\infty}(T')^2 dz = -\int_{T_S^*}^{T_F^*} f(T,P)dT = \int_{T_o}^{T_S^*} f(T,P)dT - \int_{T_o}^{T_F^*} f(T,P)dT. \qquad \text{(B4)}$$

The RHS in the above equation is the potential function U. Thus, the velocity of the savanna-forest boundary is given by

$$v = c[\mathrm{U}(T_F^*,P) - \mathrm{U}(T_S^*,P)] = c\Delta\mathrm{U}(P), \qquad \text{(B5)}$$

where $\Delta\mathrm{U}(P)$ is the difference between the values of the potential function corresponding to forest $[\mathrm{U}(T_F^*,P)]$ and savanna $[\mathrm{U}(T_S^*,P)]$ state, respectively. Further, using $f(T) = (T - a_1)(T - a_2)(T - a_3)$ one can show $c = \frac{1}{\int_{-\infty}^{\infty}(T')^2 dz} \propto \sqrt{D}$ (Murray 2001), to yield

$$v \propto \sqrt{D}\,\Delta\mathrm{U}(P). \qquad \text{(B6)}$$

#### B.2 Curvature effects in a two-dimensional landscape

To derive the expression for curvature effects in equation (7), we follow the same procedure described in the above section for a 1D model. However, we make two small changes: first, we use the polar representation of the Laplacian operator $\left[\frac{\partial^2}{\partial r^2} + \frac{1}{r}\left(\frac{\partial}{\partial r}\right) + \frac{1}{r^2}\left(\frac{\partial}{\partial \theta^2}\right)\right]$; second, we use a two-dimensional waveform of $T$, $T(r,t) = T(r - vt)$. Substituting $T(r - vt)$ in equation (2) with a 2D Laplacian we get (see the calculation for the 1D model in Appendix B.1)

$$\Delta\mathrm{U}(P) = v\int_{-\infty}^{\infty}(T')^2 dz + \int_{-\infty}^{\infty}\frac{D}{r}(T')^2 dz. \qquad \text{(B7)}$$

It should be noted that we ignored $\frac{1}{r^2}\left(\frac{\partial^2}{\partial \theta^2}\right)$ term in the 2D Laplacian operator due to local angular symmetry (Merriman et al. 1992). Assuming that the width of the boundary is much smaller as compared to the size of the landscape (see Fig. A1), we can take out $D/r$ term outside the second integral, since $(T')^2$ is always zero except at the boundary (Fig. A1), to yield

$$\Delta \mathrm{U}(P) = (v + \frac{D}{r_{Mc}}) \int_{-\infty}^{\infty} (T')^2 dz, \tag{B8}$$

where $r_{Mc}$ is the local radius of curvature of the savanna-forest boundary (Allen and Cahn 1979; Chen 1992; Evans et al. 1992; Merriman et al. 1992) (Fig. A2). For practical purposes, the radius of curvature of the boundary is approximately equal to the radius of curvature of the Maxwell precipitation contour (to be discussed later in detail). The equilibrium position of the boundary under a precipitation gradient can be calculated by setting $v = 0$, yielding

$$\Delta \mathrm{U}(P) = \frac{\sqrt{D}}{r_{Mc}}. \tag{B9}$$

Taylor expanding $\Delta \mathrm{U}$ around $P_M$, we get

$$|\Delta P| \propto \sqrt{D}\, |\kappa_{Mc}|, \tag{B10}$$

where $\Delta P$ is the local deviation of the boundary from the Maxwell precipitation contour ($P_{Mc}$), measured in units of precipitation, and $|\kappa_{Mc}|$ is the absolute curvature value of the savanna-forest boundary (Fig. A2).

In our calculations above, we assume that the curvature of the savanna-forest boundary is equal to the curvature of $P_{Mc}$. Strictly speaking, this assumption is incorrect as the curvature of the boundary is always less than or equal to the curvature of the $P_{Mc}$. However, the simulations show that this is a reasonable approximation of the actual dynamics. For example, in the simulations with linear precipitation contour (Fig. 3A), the boundary exactly coincides with the $P_{Mc}$. In this case, both the boundary and $P_{Mc}$, have the same local curvature ($|\kappa_{Mc}| = 0$). But, if $P_{Mc}$ becomes slightly curved (Fig. 3B), initially the boundary rearranges itself to track $P_{Mc}$ (Eq. 6), followed by a small deviation to account for the curvature effects in equation (7). But the change in the curvature of the boundary due to rearrangement is much greater than the change in curvature due to curvature effects. Therefore, it is reasonable to assume that the curvature of the boundary is approximately equal to the curvature of $P_{Mc}$.

### Appendix C: Integro-differential equations

In the main text, we used a reaction-diffusion equation to model spatial dynamics of savanna-forest ecosystems. However, this modeling approach is based on the assumption that seeds are dispersed locally, ignoring the effects of long-range seed dispersal. Studies on species invasion (Kot et al. 1996) and infectious diseases (Medlock and Kot 2003) show that the inclusion of long-range dispersal can qualitatively change results. For example, long-range dispersal of organisms can result in an accelerating expansion, instead of the constant rate of expansion predicted by the reaction-diffusion model (Eq. 4). Here, we show that, in contrast to past work (Kot et al. 1996; Medlock and Kot 2003), long-range spatial interactions in a bistable system have no effect the model dynamics and yield same results as the reaction-diffusion model (Barton and Turelli 2011). The reason for this, in plain terms, is that a small number of seeds (which is what happens with long-range dispersal) to a site far from the boundary are not sufficient to flip that site to the other stable state, such that long-range events do not appreciably affect the dynamics of the spatial systems.

To investigate the role of long-range seed dispersal on the dynamics of savanna-forest ecosystems we use an integro-differential equation

$$\frac{\partial T(x,t)}{\partial t} = f(T,P) + \int_{-\infty}^{\infty} K(|x' - x|)[T(x',t) - T(x,t)]dx'. \quad \text{(C1)}$$

This equation offers a simple and generic mathematical formalism to incorporate long-range spatial interactions between individuals via local interactions and dispersal. In the above equation, the spatial interactions are modeled using a convolution operator (integral) that describes the interaction of trees at a given patch with other adjacent patches. The convolution operator consists of a kernel density function, $K(x)$, which describes the contribution of spatial interactions to the growth rate (Clobert 2012). The convolution term increases as the distance between the patches decreases and the difference between their relative tree abundance increases. Therefore, spatial interactions are most pronounced at the savanna-forest boundary, where the adjacent patches of vegetation have a high relative difference in abundance. In this paper, we use symmetric kernel functions, $K(x) = K(-x)$, to model directionally unbiased spatial interactions. Furthermore, the kernel function is normalized to unity, $\int_{-\infty}^{\infty} K(x)dx = 1$. It can be easily shown that if we plug a Gaussian kernel (which assumes local spatial interactions)

$$K(x) = \frac{1}{\sqrt{\pi}\alpha} e^{-\frac{x^2}{\alpha^2}} \quad \text{(C2)}$$

into equation (C1), we recover the reaction-diffusion equation. Let $\Delta x = x - x'$. Taylor expand $T(x',t)$ about $x$, to get

$$\frac{\partial T}{\partial t} = f(T,P) + \int_{-\infty}^{\infty} \frac{1}{\sqrt{\pi}\alpha} e^{-\frac{x^2}{\alpha^2}} [\frac{\partial T}{\partial x}\Delta x + \frac{\partial^2 T}{\partial x^2}(\Delta x)^2] d(\Delta x). \quad \text{(C3)}$$

The first term in the above integral vanishes since $K(x)$ is a symmetric function, to yield a reaction-diffusion equation

$$\frac{\partial T}{\partial t} = f(T,P) + D\,\nabla^2 T, \quad \text{(C4)}$$

where $D(= \alpha^2/4)$ is the diffusion coefficient.

Next, we simulate the distribution of savanna and forest for various dispersal kernels with varying degree of fatness in their tail, keeping their mean dispersal distance (length scales of spatial interactions) constant (table C1). We find that our simulation results for each of these kernels are qualitatively the same as those obtained from the reaction-diffusion equation (Fig. C1). In other words, the speed of the savanna-forest boundary and curvature dynamics follow equations (4) and (7), respectively. These simulation results are also supported by analytical work of Bates et al. (1997), who showed that in a one-dimensional landscape for any generalized kernel (including a Gaussian kernel) the wavefront (boundary) moves with a constant speed given by equation (4), provided $\int_0^\infty |x|K(x)dx \neq \infty$.

These results at first might seem contradictory to previous studies on species invasion. To resolve this contradiction we need to understand the mechanism by which a traveling-wave is formed. In species invasion literature (Kot et al. 1996), the authors present their analysis for 'logistic' type models (stable state invading an unstable state), where even occasional long-dispersal events can have significant effects on the speed of propagation due to a positive growth rate at low densities far away from the wavefront. Since the tail of the dispersal kernel plays a key role in the propagation of the fronts, these fronts are commonly referred to as 'pulled fronts.' However, in bistable systems [one stable state invading another stable state, like the savanna-forest system described in the main text (Staver et al. 2011; Staver and Levin 2012) or an Allee effect (Keitt et al. 2001)], random long-dispersal events do not affect the speed of propagation due to negative growth rate at low densities. In these systems, the front propagates due to the dispersal of individuals from the bulk of the population to the low-density areas near the boundary. Therefore, these fronts are commonly referred to as 'pushed fronts.' We refer our readers to Van Saarloos (2003) for an extensive review on properties of front propagation.

To summarize, we show that long-range spatial interactions have no or little effect on the boundary dynamics of savanna-forest ecosystems.

**Table C1.** List of spatial kernels and the corresponding parameter values used to simulate spatial distribution of savanna and forest in a 2D landscape in figure C1. In this paper, we used four different kernels to simulate the distribution of savanna and forest (column 1 and 2). All the kernels are symmetric and have the same mean dispersal distance. But these kernels vary in the degree of the flatness of their tail (key: VTT — very thin-tail; TT — thin-tail; FT — fat-tail; VFT — very fat-tail ). Length scales for these kernels (column 3) were calculated by evaluating $\int_0^\infty 2\pi r^2 K(r,\alpha) dr$ (except for Cauchy kernel, as the integral is not defined for a Cauchy distribution). We chose the parameter value of $\alpha$ such that the length scales for all kernels remain same (column 4 and 5).

| **Kernel** | **Function** $K(r,\alpha)/K(x,y,\alpha)$ | **Length scales (formula)** | **Length scales (default value)** | **Parameter (value $\alpha$)** |
|---|---|---|---|---|
| Gaussian (VTT) | $\frac{1}{\pi\alpha^2} e^{-(\frac{r}{\alpha})^2}$ | $\frac{\alpha\sqrt{\pi}}{2}$ | 10 | $\frac{20}{\sqrt{\pi}}$ |
| Exponential (TT) | $\frac{1}{2\pi\alpha^2} e^{-(\frac{r}{\alpha})}$ | $2\alpha$ | 10 | 5 |
| Root Exponential (FT) | $\frac{1}{24\pi\alpha^2} e^{-(\frac{r}{\alpha})^{1/2}}$ | $20\alpha$ | 10 | 0.5 |
| Cauchy (VFT) | $\frac{\alpha}{2\pi} \frac{1}{(x^2+y^2+\alpha^2)^{3/2}}$ | *NA* | *NA* | 3 |

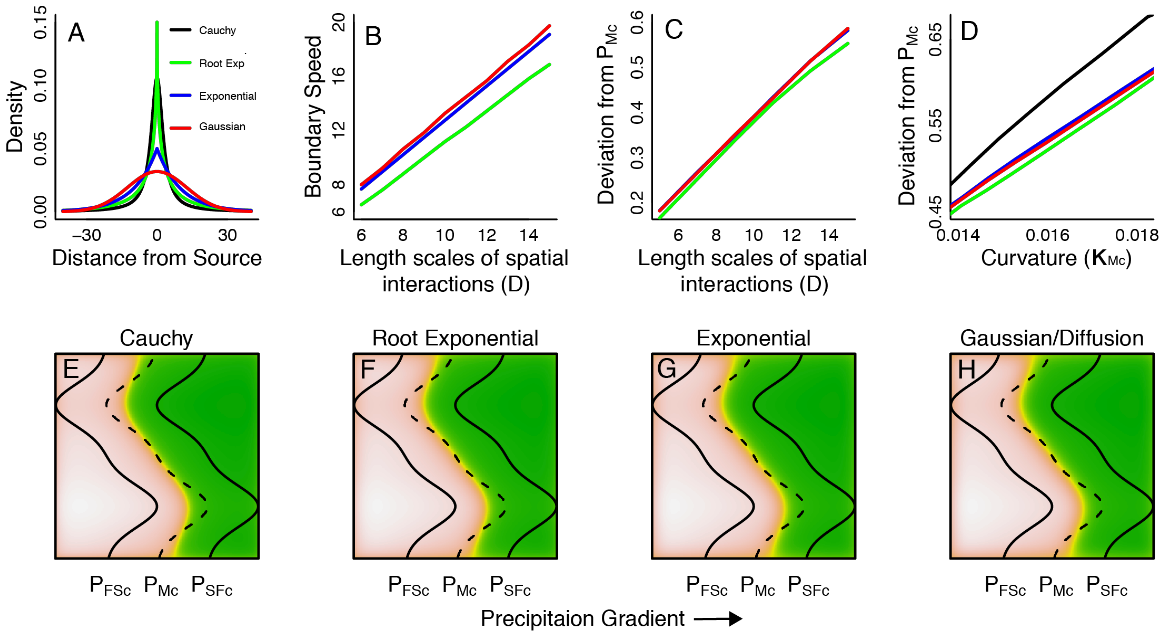


**Figure C1.** Simulations show that the boundary features remain qualitatively similar with long-range kernels. This figure is based on simulation results of the generalized 2D spatial kernels in a simple catastrophic bifurcation model (Eq. 1). Spatial distribution of savanna and forest with long range (E-F) and short-range spatial interactions (G-H) (see table C1). For all kernels in the plot (B), the speed of the boundary varies linearly with the length scales of spatial interactions (as predicted by Eq. 4). Furthermore, the deviation of the savanna-forest boundary from the $P_{Mc}$ is also linearly proportional to the length scales of spatial interactions (C) and $\kappa_{Mc}$ (D) (as predicted by Eq. 7). (E-H) The qualitative dynamics of savanna and forest distribution is independent of the choice of spatial kernel. For a generalized spatial kernel, the length scales were calculated by evaluating $\int_0^\infty 2\pi r^2 K(r,\alpha)dr$ (see table C1). In plots (D-H) we chose the values of $\alpha$ such that the length scales for all kernels remain same, except Cauchy, since the above integral is not finite for a Cauchy distribution. These simulations were performed on a lattice of size $256$ x $256$ with $D = 10$ (see Numerical Methods in Online Appendix D).

**Online Appendix D: Supplementary Methods and Figures**

**Part A: Models**

In this paper, we use four catastrophic bifurcations models to study the role of space in alternative stable states ecosystems. The first model (Eq. 1) is a simple savanna-forest model developed by Staver and Levin (2012) (see main text for details). Models, two (Eq. D1) and three (Eq. D2) are single-variable dynamical models for which we can define a potential function (Strogatz 2014). The fourth model (Eqs. D3 and D4) is a two-variable coupled dynamical system with no defined potential function (Van Geest et al. 2007). We choose these models only for numerical simulations. Analytical results presented in the main text are robust to the choice of model.

*Model 2: A simple catastrophic bifurcation model*

The second model is an imperfect catastrophic bifurcation model with the following dynamical equation (Strogatz 2014)

$$\frac{dT}{dt} = -P + rT - T^3 = f(T, P) \qquad \text{(D1)}$$

where $T$ is the state of the system (tree cover), and $P$ is the control parameter (precipitation). The roots of this equation are obtained by setting $f(T^*, P) = 0$, where $T^*$ corresponds to roots of the system for a given value of control parameter. We use $T_S^*$ and $T_F^*$ to denote stable roots of the system, corresponding to savanna and forest state, respectively. The roots are stable (unstable) if $df/dT|_{T=T^*} < 0$ ($> 0$). Furthermore, the critical precipitation ($P_{FS}$ and $P_{SF}$) values of the system are obtained by simultaneously solving $f(T^*) = 0$ and $df/dT|_{T=T^*} = 0$. $P_{FS}$ (or $P_{SF}$) $= \pm 2(r/3)^{\frac{3}{2}}$. We obtained the Maxwell precipitation ($P_M$) by setting $\int_{T_S^*}^{T_F^*} f(T, P) = 0$. In this paper, we use $r = 3$ as the default parameter value. We use this model for main text figures 5, 6D and D4.

Using the mathematical conditions above, we obtained $P_{FS} = -2$, $P_{SF} = 2$, and $P_M = 0$. Please note that we only use relative values of precipitation and vegetation. Positive values of vegetation and precipitation correspond to the forest and higher precipitation regions, respectively. Similarly, negative values of vegetation and precipitation correspond to savanna and lower precipitation regions, respectively.

*Model 3: Harvesting Model*

$$\frac{dV}{dt} = rV\left(1 - \frac{V}{K}\right) - c\frac{V^2}{V_o^2 + V^2} \qquad \text{(D2)}$$

The third model we use in this paper is an insect outbreak population model used by Ludwig to analyze spruce budworm outbreak in eastern Canada (Ludwig et al. 1978; May 1977). This model has also been used to study vegetation dynamics (Guttal and Jayaprakash 2007; Guttal and Jayaprakash 2008; Noy-Meir 1975; van de Leemput et al. 2015) in semiarid ecosystems and fishery dynamics (May 1977). Here, $V$ represents the state of the system, e.g., biomass or spruce density. The model consists of two terms. The first term is a logistic growth function with

$r$ and $K$ as the per-capita growth rate for small population sizes and carrying capacity of the system, respectively. The second term is the harvesting term with $c$ as the harvesting rate (control parameter) and $V_o$ as the half-saturation constant. In this paper, we use $r = 1$, $K = 10$, and $V_o = 1$ as default parameter values.

Using the above parameter values, we calculated the critical harvesting parameter to be $c_1 = 1.8$ and $c_2 = 2.6$. Furthermore, we found $c_M = 2.35$. We used this model to study the dynamics of the system under additive and multiplicative noise (Fig. D1)

*Model 4: Light attenuation model*

$$\frac{dV}{dt} = r_v V(1 - V\frac{h_E^P}{E^p}) \qquad \text{(D3)}$$

$$\frac{dE}{dt} = r_E E(1 - \frac{E}{E_o}\frac{h_v + V}{h^v}) \qquad \text{(D4)}$$

Finally, we consider a two-variable macrophyte and turbidity model. This model has been used to describe lake-eutrophication in shallow lake water ecosystem (Van Geest et al. 2007). The two dynamical variable V and E are macrophyte and vertical light attenuation, respectively, with $E_o$ as the control parameter of the system. The control parameter describes the vertical light attenuation in the absence of macrophytes. $E_o$ is a proxy used in place of the nutrient loading in the lake. See Scheffer (2004) for further details on the model.

We use $h_E = 2$, $h_V = 0.2$, $P = 4$, $r_E = 0.1$ and $r_V = 0.05$ as default parameter values. Critical points of this model are calculated by setting the determinant of the Jacobian to zero (Strogatz 2014) ($E_{o1} = 5.2$ and $E_{02} = 7.3$). This model has no potential function, and the Maxwell point is dependent on the ratio of the diffusion coefficients ($D_v/D_e$; Fig. D2) (van de Leemput et al. 2015).

**Part B: Numerical Methods**

All the numerical simulations were performed using R (version 3.2.2) and MATLAB (R 2015a). By default, in all the simulations, we used reflective boundary conditions and initialed them with random numbers from standard uniform distribution. These simulation methods closely follow the methods described in Guttal and Jayaprakash (2009).

**A Numerical scheme for simulating reaction-diffusion in a 1D landscape**

We performed numerical simulations for the 1-D reaction-diffusion model using the following Euler forward time scheme that is first-order accurate in time and second order accurate in space:

$$T_i^{t+1} = T_i^t + \Delta t[f(T_i^t, P_i) + D(T_{i-1}^t + T_{i+1}^t - 2T_i^t)], \qquad \text{(D5)}$$

where i represents the cell coordinate and t represents the time step. The simulations were truncated when $\sum_i |T_i^{t+1} - T_i^t| < 0.1$ because near equilibrium the change in tree density over successive time steps is very small. This numerical scheme was used to simulate figure 2

**A Numerical scheme for simulating reaction-diffusion in a 2D landscape**

*Reaction-diffusion equation without noise*

For 2-D reaction-diffusion model, using the following Euler forward time scheme that is first order accurate in time and second order accurate in space:

$$T_{i,j}^{t+1} = T_{i,j}^{t} + \Delta t[f(T_{i,j}^{t}, P_{i,j}) + D(T_{i-1,j}^{t} + T_{i+1,j}^{t} + T_{i,j-1}^{t} + T_{i,j+1}^{t} - 4T_{i,j}^{t})], \quad \text{(D6)}$$

where $i$ and $j$ represent the cell coordinates. Similar to the 1-D model, we truncated the simulations when $\sum_{i,j} |T_{i,j}^{t+1} - T_{i,j}^{t}| < 0.1$. This numerical scheme was used to simulate figures 3, 5 and D2.

*Reaction-diffusion equation with noise*

$$\frac{\partial T}{\partial t} = f(T, P) + D\,\nabla^2 T + \sigma g(T, P)\eta(x, y, t) \quad \text{(D7)}$$

To simulate the above stochastic reaction-diffusion equation, we used the following numerical scheme:

$$T_{i,j}^{t+1} = T_{i,j}^{t} + \Delta t[f(T_{i,j}^{t}, P_{i,j}) + D(T_{i-1,j}^{t} + T_{i+1,j}^{t} + T_{i,j-1}^{t} + T_{i,j+1}^{t} - 4T_{i,j}^{t})] + \sigma\sqrt{\Delta t}g(T_{i,j}^{t})\eta_{i,j}^{t} \text{(D8)}$$

where $\eta_{i,j}^{t}$ is a standard normal random variable and $\sigma$ is the standard deviation. This numerical scheme was used to simulate figure D1.

*Integro-differential equations*

For a generalized kernel in equation (1) we used the following numerical scheme:

$$T_{i,j}^{t+1} = T_{i,j}^{t} + \Delta t(f(T_{i,j}^{t}, P_{i,j}) + D\sum_{i',j'} K(i - i', j - j')[T_{i,j}^{t} - T_{i',j'}^{t}]) \quad \text{(D9)}$$

To reduce the computational time, we only evaluated the convolution integral (summation) for $95\%$ confidence interval and then re-normalized the kernel to unity. The convolution was solved using *conv2* function in MATLAB. The simulations were truncated when $\sum_{i,j} |T_{i,j}^{t+1} - T_{i,j}^{t}| < 0.1$. This numerical scheme was used to simulate figure C1.

**Part C: Data Analysis (Methods)**

All the data analyses were performed using R 3.2.2.

**Data Sources**

*Precipitation:*

We used monthly precipitation records derived from Tropical Rainfall Measuring Mission monthly precipitation product [TRMM 3B43; see Huffman and Bolvin (2013) for documentation]. To get a mean annual estimate for precipitation, we averaged the precipitation over a period of $10$ years (January 1, 2000 to December 31, 2009), for Sub Saharan Africa ($20.5^o\ W - 53.75^o\ E$ and $20^o\ S - 21^o\ N$). The TRMM dataset is prepared using a combination of satellite data, precipitation gauges and modeled precipitation based on cloud cover. To estimate curvature of the savanna-forest boundary we projected the original precipitation raster of $0.25^o$ x $0.25^o$ to a Lambert equal area precipitation raster (using *projectRaster* function) of resolution $25$ km x $25$ km (henceforth referred as low-resolution precipitation raster). For large-scale spatial simulations, we increased the resolution of the original precipitation raster ($0.25^o$ x $0.25^o$) by a factor of $4$ (using a *resample* function with bilinear interpolation). We then projected it to a Lambert equal-area projection (using *projectRaster* function), to get a precipitation raster of resolution $12.5$ km x $12.5$ km (henceforth referred as high-resolution precipitation raster). This was done because the length scales of spatial interactions were the order of $25$ km (preliminary simulation results using low-resolution raster), i.e., on the same scale as the dimensions of a single cell. This prevented the savanna-forest boundary from reaching its equilibrium position due to pinning (Keitt et al. 2001; see also our additional forthcoming work). All the cells corresponding to water bodies (like lakes and oceans) were omitted.

*Vegetation:*
We used a freely available global tree cover distribution product produced by Hansen et al. (2013) using Landsat, which reports relative tree cover distribution ranging from values between $0\%$ (completely open canopy) to $100\%$ (completely closed canopy). We chose the minima between the two modes (savanna and forest) of tree cover as the savanna-forest boundary. Using this procedure, we identified $76\%$ as the threshold tree cover ($B_v$) for the boundary, although our results were robust to the slight variations to this threshold (Fig. D3). The vegetation raster was initially made to the same resolution of original precipitation raster ($0.25^o$ x $0.25^o$). Then we followed the same steps described in the preparation of precipitation raster to get a low- and high-resolution vegetation raster.

**Estimating Maxwell precipitation from empirical data**

To estimate the curvature of the savanna-forest boundary, we used the *contourLines* function to extract the cell coordinates of the boundary ($B_v$) from the low-resolution vegetation raster (a total of $N$ pair of $x$ and $y$ cell coordinates). These coordinates were real values and not integers. Since the typical length of the boundary was the order of $10^3 - 10^4$ km. All the boundaries of length less than $150$ km were ignored, as they were too small to estimate curvature. Then, for each boundary, we took a rolling window of segment length $2n + 1$ points (a pair of $x$ and $y$ cell coordinates) and estimated the radius of curvature (or curvature, the inverse of the radius of curvature) by fitting a circle to the cell coordinates of the segment using *lsfit.circle* function. The cell coordinates were then transformed to actual distances (1 unit distance in cell coordinates = $25$ km). To estimate the precipitation corresponding to this curvature value, we took the precipitation value corresponding to the midpoint of the segment ($(n + 1)^{th}$ point). Since the midpoint was not an integer, we rounded midpoints to closest integer values so that precipitation values could be extracted from the low-resolution precipitation raster. We then moved the rolling window by $1$ point and repeated the same procedure. For the $n$ points at the beginning ($1$ to $n$) and end ($N + 1 - n$ to $N$) of the boundary for which the curvature could not be estimated, we used the curvature values at $(n + 1)^{th}$ and $(N - n)^{th}$ points, respectively.

To estimate $P_M$, we fitted equation 7 to the curvature and boundary precipitation data using *nls2* function (black V-shape curve in fig 4C). For better visualization, we grouped precipitation values into $11$ precipitation intervals of $200$ mm each, ranging from $800$ mm to $3000$ mm. For each of these precipitation intervals, we grouped the corresponding curvature values into one category and plotted the mean values of the curvature in each category against the mid-point of each precipitation interval (like $900$ mm, $11000$ mm and so on). We also estimated $95\%$ confidence interval for curvature values in each precipitation category (grey spindles in Fig. 4)

In this paper, we provide a simple method to estimate $P_M$ by exploiting the fact that the boundaries deviate linearly from the Maxwell contour as a function of the local curvature of the boundary (also see large-scale simulations below for an alternate method to estimate $P_M$). We argue that this method of estimating $P_M$ improves upon the method proposed by Staal et al. (2016), which uses the concept of potential landscapes, for two reasons. First, their methods use only one-dimension to model space, which provides an incomplete picture of the actual dynamics of savanna and forest ecosystems. Here, we show that including a second dimension qualitatively changes the dynamics of the system. Second, they estimate $P_M$ by reconstructing the potential function of the system using a mean-field approach. If one assumes that savanna-forest systems show strong spatial interactions, the potential function of the system is modified to (Malchow et al. 1983; Schlögl 1972)

$$\mathrm{U} = \int \left(\frac{D}{2}(\nabla T)^2 - \int f(T,P)dT\right)dr, \tag{D10}$$

and one can no longer use the mean-field potential function. Nevertheless, mean-field potential function provides valuable insights (see main text) and should be used, but with caution.

**Sensitivity Analysis**

For estimating $P_M$, we used two parameters: the vegetation cover corresponding to the savanna-forest boundary ($B_v$) and the length of boundary segment (rolling window) for which the curvature values were estimated ($n$). To check whether our estimate of $P_M$ was sensitive to specific values of the parameter, we computed $P_M$ for $8$ values of $B_v$ (from $73$ to $80$ in steps of $1$) and $20$ values of $n$ (from $6$ to $25$ in steps of $1$). The sensitivity analysis showed that the estimate of $P_M$ was robust to the choice parameters ($B_v$ and $n$) and lay within the range of $1508 \pm 84$mm MAP, with $1508$ mm MAP and $1513$ mm MAP as the median and mode of $P_M$ estimates, respectively (Fig. D3).

**Large-scale simulations**

The model results indicate that the equilibrium position of the boundary is dependent on two independent parameters: $P_M$ (Eq. 4), and the product of $\kappa_{Mc}$ and $D$ (Eq. 7). For a landscape with linear precipitation contours, the boundary always aligns with $P_{Mc}$ irrespective of the choice of $D$ and the underlying model (Fig. 3A). However, if the precipitation contours are curved, the boundary deviates from $P_{Mc}$, and these deviations are proportional to the product of the diffusion coefficient and a model dependent constant (Fig. 3B and Eq. 7). Exploiting this feature of the bistable reaction-diffusion model, we argue that the choice of the model will not affect the estimate of $P_M$, but it may not give us accurate length scales of spatial interactions.

We use imperfect catastrophic bifurcation model in our large-scale spatial simulations. For a given value of $P_M$, we rescale the high-resolution precipitation raster ($P_{old}$) by the following relation

$$P_{new} = (P_{old} - P_{Mc})/250. \qquad \text{(D11)}$$

We use this to rescale precipitation raster because we know from previous studies that the critical points corresponding to forest to savanna and savanna to forest transition are around 1000 mm and 2000-2600mm, respectively (Aleman and Staver 2018; Staver et al. 2011). Furthermore, the curvature calculations indicate that $P_M$ is around 1500mm (see the estimation of $P_M$ from curvature and precipitation data). If we plug these estimates of critical values and Maxwell point in the above relationship (Eq. D11), we recover back the critical values and Maxwell point corresponding imperfect catastrophic bifurcation model (Eq. D1).

Next, we rescale the high-resolution vegetation raster ($T_{old}$) by the following relation

$$T_{new} = (T_{old} - B_v). \qquad \text{(D12)}$$

In these simulations, we use $B_v = 76\%$. We use this to rescale vegetation raster because we know that the imperfect bifurcation model has a boundary at $T = 0$. In this rescaled vegetation raster ($T_{new}$), all the cells above (below) zero are forest (savanna). For a given value of $P_M$ and $D$, we numerically simulate the distribution of savanna and forest using the rescaled precipitation raster ($P_{new}$) as the input (see numerical simulations). All the cells corresponding to water bodies in the simulated vegetation raster ($T_{sim}$) were reclassified as zero at every time step of the simulation since they neither act as savanna or forest.

Now we define a fitness function $F$ such that $F =$ (number of forest cells in $T_{sim}$ that are also forest in $T_{new}$ $+$ number of savanna cells in $T_{sim}$ that are also savanna in $T_{new}$) $-$ (number of forest cells in $T_{sim}$ that are also savanna in $T_{new}$ $+$ number of savanna cells in $T_{sim}$ that are also forest in $T_{new}$).

In other words, the fitness function is maximum when the simulated distribution of forest and savanna exactly match the empirical data and is minimum when there is a complete mismatch. The simulated vegetation matrix $T_{sim}$ is initialized with $T_{new}$ (current distribution of savanna and forest).

Next, we use a Genetic Algorithm (Scrucca 2013) to find parameter values of $P_M$ and $D$ that maximize this fitness function. In the GA simulation, we used the following default parameter values: type = "real-valued", $D_{min} = 4$, $D_{max} = 100$, $(P_M)_{min} = 1400$ mm, $(P_M)_{max} = 1600$ mm, popSize $= 10$, Maxiter $= 50$ and parallel=T.

We excluded all the areas in Western Africa and edaphic savannas of Bateke plateau from our analyses. Next, using the optimized parameter values of $P_M$ and $D$, we simulated the distribution of savanna and forest for two other initial conditions. For the first simulation, we initialized all the cells as savanna (Fig. 5B). In the second simulation, we initialized all the cells as forest (Fig. 5C). See main text for interpretation of these results.

**Model comparison**

To compare the performance of various biome distribution model, we simulated biome patterns in Sub Saharan Africa using three biome distribution models: (*a*) 'one-climate one-biome' model (Fig. 6A); (*b*) mean-field bistable model (Fig. 6B); and (*c*) 2D reaction-diffusion model (Fig. 6C). In these simulations, we excluded the contribution of the deforested regions in Western Africa and the Bateke Plateau from the fitness function (see Data Analysis in Online Appendix D).

*'One-climate one biome' model:* We assumed a unique precipitation contour that is coincident with the savanna-forest boundary. Below (above) this precipitation value the equilibrium biome state in the landscape is savanna (forest). We estimated this precipitation value by using a Genetic Algorithm (maximizing the fitness function) described in the Large-scale simulations section.

*Mean-field bistable model:* For this model, we simulated biome patterns using the imperfect catastrophic bifurcation model (see Models in Online Appendix D) with rescaled precipitation as input data (Eq. D11). In this model, we initialized the simulation with randomly distributed savanna and forest patches.

*2D reaction-diffusion model with bistable reaction part:* We followed the same procedure as described in the Large-scale simulations section.

**Part D: Supplementary Figures**

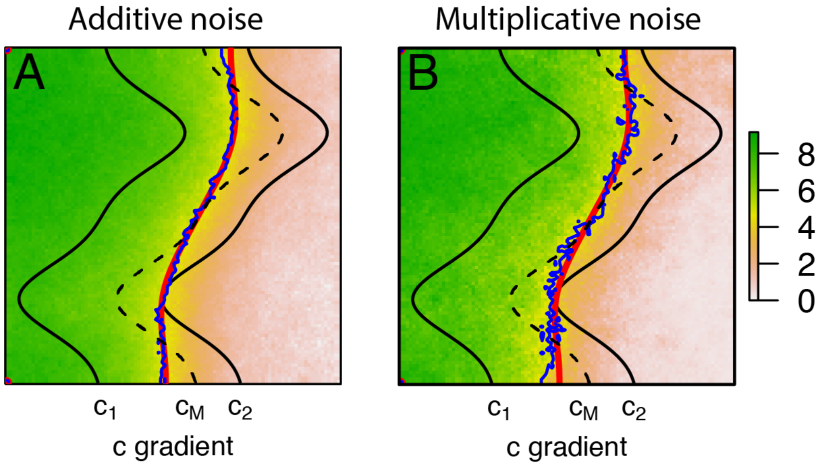


**Figure D1:** Savanna-forest boundary dynamics with (A) additive noise, $g(x) = 1$, and (B) multiplicative noise, $g(x) = N^2/(N_o^2 + N^2)$ . The blue and red line in plots represents the position of the boundary with and without the noise term, respectively. In both plots (A) and (B), we see that the addition of noise does not change the qualitative behavior of the system. Simulations were truncated after $80$ time units. These simulations were performed using Harvesting model (see Models in Online Appendix D) on a $100\text{x}100$ lattice. Here, we used $D = 50$ and $\sigma = 2$ (see Numerical Methods in Online Appendix D).

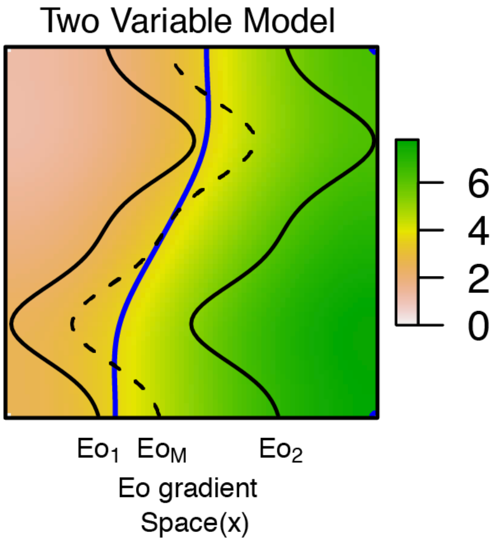


**Figure D2:** The simulation results form the two-variable light attenuation model (see Models in Online Appendix D) with no potential function is qualitatively similar to the one variable model. However, unlike the one variable model, the equilibrium position of the boundary between the two stable states is also dependent on $D_v/D_E$. These simulations were performed on a $100\text{x}100$ lattice. Here, we used $D_E = D_v = 10$ (see Numerical Methods in Online Appendix D).

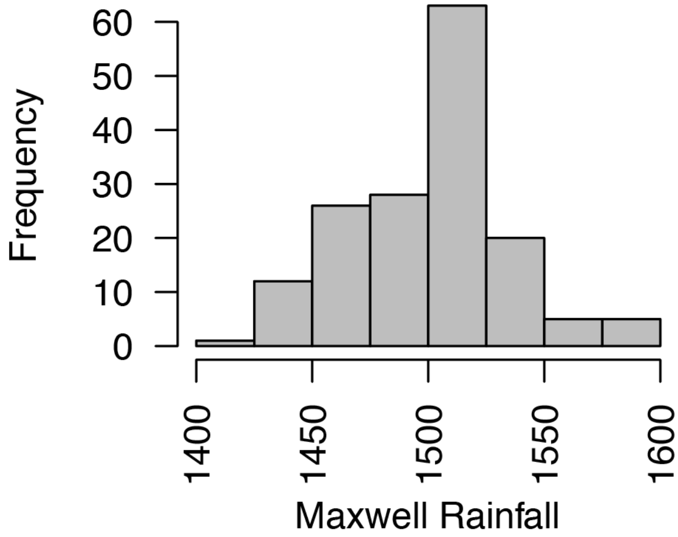


**Figure D3:** Sensitivity histogram of $P_M$ estimated for various combinations of parameter values (Boundary Vegetation $B_v$ and length of boundary segment used to calculate curvature) used in data analysis. The histogram suggests that the estimated value of $P_M$ is robust to the changes in the parameter values and lies within $1508 \pm 84$ mm MAP, with $1508$ mm MAP and $1513$ mm MAP as the median and mode of $P_M$ estimates (see Data Analysis in Online Appendix D)

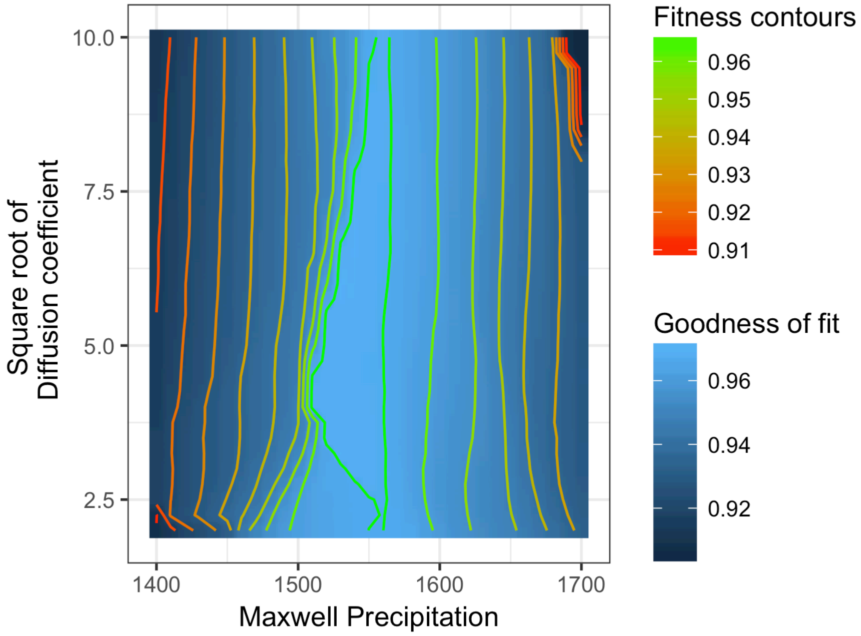


**Figure D4:** Comparing the simulated distribution of biomes (using a 2D reaction-diffusion model) with the current distribution of biomes in Sub Saharan Africa for various combinations of parameter values of $P_M$ and $\sqrt{D}$. In these simulations, we excluded the contribution of the deforested regions in Western Africa and edaphic savannas of Bateke Plateau from the fitness function. The sensitivity analysis suggests that distribution of biomes in Sub Saharan Africa is relatively independent of $D$. Furthermore, the finesse function is maximum near $P_M = 1500 -$ $1550$ mm MAP, which is consistent with the estimate of $P_M$ obtained from the curvature analysis in figure 4C (see Data Analysis in Online Appendix D).

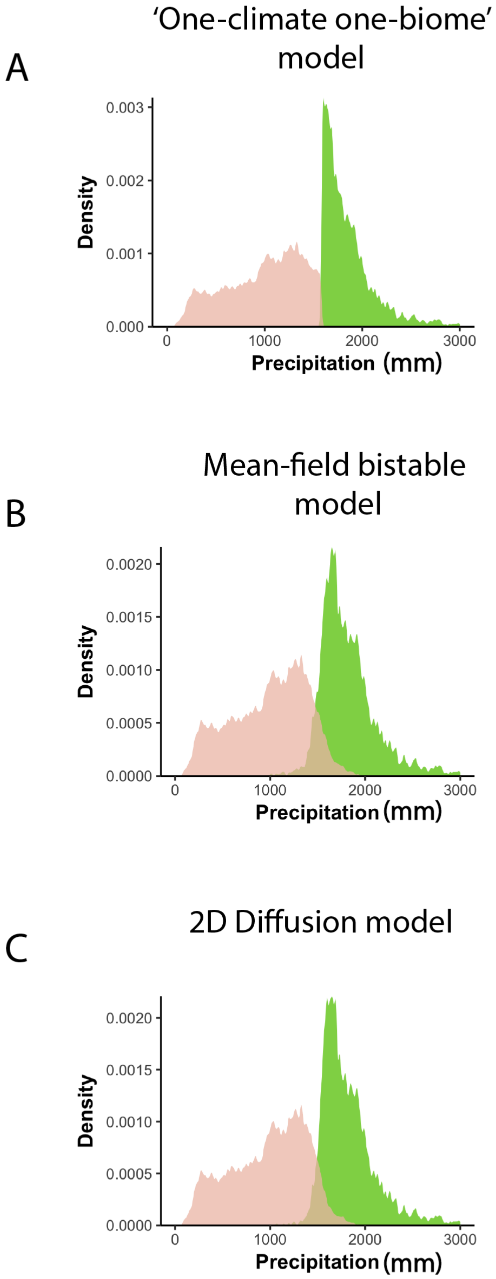


**Figure D5:** Simulated climatic niches of savanna and forest biomes from three biome distribution models: (A) mean-field bistable model, (B) classical biome distribution model, and (C) 2D reaction-diffusion model with a bistable reaction part (see Data Analysis in Online Appendix D). Our simulations suggest that all models, except the classical biome distribution model, can generate empirically observed overlap in rainfall ranges of savanna and forest biomes.

### Refrences


Aleman, J. C., and C. A. Staver. 2018. Spatial patterns in the global distributions of savanna and forest. Global Ecology and Biogeography 0.

Allen, S. M., and J. W. Cahn. 1979. A microscopic theory for antiphase boundary motion and its application to antiphase domain coarsening. Acta Metallurgica 27:1085--1095.

Aronson, D. G., and H. F. Weinberger. 1975. Nonlinear diffusion in population genetics, combustion, and nerve pulse propagation, Pages 5–49 Partial differential equations and related topics, Springer.

Barton, N., and M. Turelli. 2011. Spatial waves of advance with bistable dynamics: cytoplasmic and genetic analogues of Allee effects. The American Naturalist 178:E48—E75.

Bates, P. W., P. C. Fife, X. Ren, and X. Wang. 1997. Traveling waves in a convolution model for phase transitions. Archive for Rational Mechanics and Analysis 138:105--136.

Bramson, M. 1983, Convergence of solutions of the Kolmogorov equation to travelling waves, v. 285, American Mathematical Soc.

Chen, X. 1992. Generation and propagation of interfaces for reaction-diffusion equations. Journal of Differential equations 96:116--141.

Clobert, J. 2012, Dispersal ecology and evolution, Oxford University Press.

Evans, L. C., H. M. Soner, and P. E. Souganidis. 1992. Phase transitions and generalized motion by mean curvature. Communications on Pure and Applied Mathematics 45:1097--1123.

Fife, P. C., and J. B. McLeod. 1977. The approach of solutions of nonlinear diffusion equations to travelling front solutions. Archive for Rational Mechanics and Analysis 65:335–361.

Guttal, V., and C. Jayaprakash. 2007. Self-organization and productivity in semi-arid ecosystems: implications of seasonality in rainfall. Journal of theoretical biology 248:490--500.

—. 2008. Changing skewness: an early warning signal of regime shifts in ecosystems. Ecology letters 11:450--460.

—. 2009. Spatial variance and spatial skewness: leading indicators of regime shifts in spatial ecological systems. Theoretical Ecology 2:3--12.

Hansen, M. C., P. V. Potapov, R. Moore, M. Hancher, S. A. Turubanova, A. Tyukavina, D. Thau et al. 2013. High-resolution global maps of 21st-century forest cover change. science 342:850--853.

Huffman, G. J., and D. T. Bolvin. 2013. TRMM and other data precipitation data set documentation. NASA, Greenbelt, USA 28.

Keitt, T. H., M. A. Lewis, and R. D. Holt. 2001. Allee effects, invasion pinning, and species? borders. The American Naturalist 157:203--216.

Kot, M., M. A. Lewis, and P. van den Driessche. 1996. Dispersal data and the spread of invading organisms. Ecology 77:2027--2042.

Ludwig, D., D. D. Jones, and C. S. Holling. 1978. Qualitative analysis of insect outbreak systems: the spruce budworm and forest. The Journal of Animal Ecology:315--332.

Malchow, H., W. Ebeling, R. Feistel, and L. Schimansky-Geier. 1983. Stochastic Bifurcations in a Bistable Reaction-Diffusion System with Neumann Boundary Conditions. Annalen der Physik 495:151--160.

May, R. M. 1977. Thresholds and breakpoints in ecosystems with a multiplicity of stable states. Nature 269:471--477.

Medlock, J., and M. Kot. 2003. Spreading disease: integro-differential equations old and new. Mathematical Biosciences 184:201--222.

Merriman, B., J. K. Bence, and S. Osher. 1992, Diffusion generated motion by mean curvature, Department of Mathematics, University of California, Los Angeles.

Murray, J. D. 2001, Mathematical Biology. II Spatial Models and Biomedical Applications ${$Interdisciplinary Applied Mathematics V. 18$}$, Springer-Verlag New York Incorporated.

Noy-Meir, I. 1975. Stability of grazing systems: an application of predator-prey graphs. The Journal of Ecology:459--481.

Scheffer, M. 2004, Ecology of shallow lakes, Springer Science & Business Media.

Schlögl, F. 1972. Chemical reaction models for non-equilibrium phase transitions. Zeitschrift für Physik A Hadrons and Nuclei 253:147--161.

Scrucca, L. 2013. GA: a package for genetic algorithms in R. Journal of Statistical Software 53:1--37.

Staal, A., S. C. Dekker, C. Xu, and E. H. van Nes. 2016. Bistability, Spatial Interaction, and the Distribution of Tropical Forests and Savannas. Ecosystems 19:1080--1091.

Staver, A. C., S. Archibald, and S. Levin. 2011. Tree cover in sub-Saharan Africa: Rainfall and fire constrain forest and savanna as alternative stable states. Ecology 92:1063--1072.

Staver, A. C., and S. A. Levin. 2012. Integrating theoretical climate and fire effects on savanna and forest systems. The American Naturalist 180:211--224.

Strogatz, S. H. 2014, Nonlinear dynamics and chaos: with applications to physics, biology, chemistry, and engineering, Westview press.

van de Leemput, I. A., E. H. van Nes, and M. Scheffer. 2015. Resilience of alternative states in spatially extended ecosystems. PloS one 10:e0116859.

Van Geest, G. J., H. Coops, M. Scheffer, and E. H. Van Nes. 2007. Long transients near the ghost of a stable state in eutrophic shallow lakes with fluctuating water levels. Ecosystems 10:37--47.

Van Saarloos, W. 2003. Front propagation into unstable states. Physics reports 386:29--222.
